# Assessing the zoonotic potential of a novel bat morbillivirus

**DOI:** 10.1101/2021.09.17.460143

**Authors:** Satoshi Ikegame, Jillian C. Carmichael, Heather Wells, Robert L. Furler O’Brien, Joshua A. Acklin, Hsin-Ping Chiu, Kasopefoluwa Y. Oguntuyo, Robert M. Cox, Aum R. Patel, Shreyas Kowdle, Christian S. Stevens, Miles Eckley, Shijun Zhan, Jean K. Lim, Ethan C. Veit, Matthew Evans, Takao Hashiguchi, Edison Durigon, Tony Schountz, Jonathan H. Epstein, Richard K. Plemper, Peter Daszak, Simon J. Anthony, Benhur Lee

## Abstract

Morbilliviruses are amongst the most contagious viral pathogens that infect mammals. Metagenomic surveys have identified numerous morbillivirus sequences in bats, but no full-length authentic morbillivirus has been isolated or characterized from bats. Here we detail the discovery of full-length Myotis Bat Morbillivirus (MBaMV) from a bat surveillance program in Brazil. After determining that MBaMV utilizes bat CD150 but not human CD150 as an entry receptor, we generated an infectious clone of MBaMV using reverse genetics. MBaMV exhibited features consistent with other morbilliviruses, including pleomorphic virions, P-editing and the rule-of-six. MBaMV replicated well in human epithelial cell lines in a nectin-4 dependent manner. Surprisingly, MBaMV was able to infect human macrophages in a CD150-independent manner. However, MBaMV was restricted by cross-neutralizing human sera and did not evade the human innate immune system, indicating that while zoonotic spillover into humans may be possible, MBaMV replication in humans would likely be restricted.

## Introduction

Bats are significant reservoir hosts for many viruses with zoonotic potential^1^. SARS-CoV-2, Ebola virus, and Nipah virus are examples of such viruses that have caused deadly epidemics and pandemics when spilled over from bats into human and animal populations^2,3^. Careful surveillance of viruses in bats is critical for identifying potential zoonotic pathogens. However, metagenomic surveys in bats often do not result in full-length viral sequences that can be used to regenerate such viruses for targeted characterization^4^, at least not without much further effort like 3’ and 5’ RACE. Improvements in sequencing technologies and bioinformatics have enabled more complete genome assemblies. Three metagenomic surveys published in the past year confirm that bats, and to a lesser extent shrews and rodents, are hosts to diverse paramyxoviruses^5–7^ that comprise multiple genera (*Jeilongvirus, Morbillivirus, Henipavirus*). Metagenomic sequences, however complete, cannot at present yield sufficiently accurate information about viral phenotypes in vitro and in vivo. Detailed virological investigations are still needed to reify taxonomic discoveries.

Morbilliviruses are amongst the most contagious viral pathogens that infect mammals. While numerous partial sequences of morbilliviruses have been identified in bats and rodents^4,5^ in metagenomic surveys, no full-length authentic morbillivirus has been isolated or characterized from chiropteran hosts. The morbillivirus genus includes measles virus (MeV), canine distemper virus (CDV), rinderpest virus, phocine distemper virus, cetacean morbillivirus, peste des petis ruminants virus and feline morbillivirus^8^. A porcine morbillivirus was recently described to be the putative cause of fetal death and encephalitis in pigs^9^. All morbilliviruses cause severe disease in their natural hosts^10–14^, and pathogenicity is largely determined by species specific expression of canonical morbillivirus receptors, CD150/SLAMF1^15^ and NECTIN4^16^.

Here, we identify and characterize a novel morbillivirus from a vespertilionid bat species (*Myotis riparius*) in Brazil, which we term myotis bat morbillivirus (MBaMV). MBaMV used *Myotis spp* CD150 much better than human and dog CD150 in fusion assays. We confirmed this using live MBaMV that was rescued by reverse genetics. Surprisingly, MBaMV replicated in primary human myeloid but not lymphoid cells and did so in a CD150-independent fashion. This is in contrast to MeV which is known to infect CD150+ human myeloid and lymphoid cells. Furthermore, MBaMV replicated in human epithelial cells and used human nectin-4 almost as well as MeV. Nonetheless, MBaMV P/V genes do not appear to antagonize human interferon induction and signaling pathways and MBaMV was cross-neutralized, albeit to variable extents, by MMR vaccinee sera. Our results demonstrate the ability of MBaMV to infect and replicate in some human cells that are critical for MeV pathogenesis and transmission. Yet comprehensive evaluation of viral characteristics provide data for proper evaluation of its zoonotic potential.

## Results

### Isolation of MBaMV sequence

During a metagenomic genomic survey of viruses in bats, we identified a full-length morbillivirus sequence from a riparian myotis bat (*Myotis riparius*) in Brazil. This myotis bat morbillivirus (MBaMV) had a genome length of 15,720 nucleotides consistent with the rule of six and comprise of six transcriptional units encoding the canonical open reading frames (ORFs) of nucleo (N) protein, phospho (P) protein, matrix (M) protein, fusion (F) protein, receptor binding protein (RBP), and large (L) protein (Extended Data Fig. 1a). The sizes of these ORFs are comparable to their counterparts in the other morbilliviruses (Extended Data Fig. 1b). Phylogenetic analysis using the full-length L protein sequence indicated that MBaMV is most closely related to canine distemper virus (CDV) and phocine distemper virus (PDV) (Extended Data Fig. 1c, Extended Data Table 1).

Paramyxovirus proteins with the most frequent and direct interactions with host proteins, such as P and its accessory gene products (V and C) as well as the RBP, tend to exhibit the greatest diversity^17^. Morbillivirus P, V and C antagonize host-specific innate immune responses while its RBP interacts with host-specific receptors. That these proteins are under evolutionary pressure to interact with different host proteins is reflected in the lower conservation of MBaMV P/V/C (31-43%) and RBP (27-32%) with other morbillivirus homologs. This is in contrast to the relatively high conservation (52-76%) of MBaMV N, M, F, and L proteins with their respective morbillivirus counterparts (Extended Data Fig. 2).

### Species specific receptor usage

The use of CD150/SLAMF1 to enter myeloid and lymphoid cells is a hallmark of morbilliviruses, and also a major determinant of pathogenicity. CD150 is highly divergent across species, and accounts for the species restricted tropism of most morbilliviruses^18^. Thus, we first characterized the species-specific receptor tropism of MBaMV. We performed a quantitative image-based fusion assay (QIFA) by co-transfecting expression vectors encoding MBaMV-F and -RBP, along with CD150 from the indicated species into receptor-negative CHO cells. MeV-RBP and F formed more syncytia in CHO cells upon human-CD150 (hCD150) co-transfection compared to dog-CD150 (dCD150) or bat-CD150 (bCD150) (Fig. 1a, top row). In contrast, MBaMV-RBP and F formed bigger and more numerous syncytia upon bCD150 overexpression than hCD150 or dCD150 (Fig. 1a, middle row). CDV-RBP and F formed extensive syncytia with both dCD150 and bCD150, and moderate syncytia with hCD150 and even mock-transfected cells (Fig. 1a, bottom row), suggesting a degree of promiscuity. We quantified these differential syncytia formation results on an image cytometer as described^19^ (Fig. 1b).

**Figure 1.**
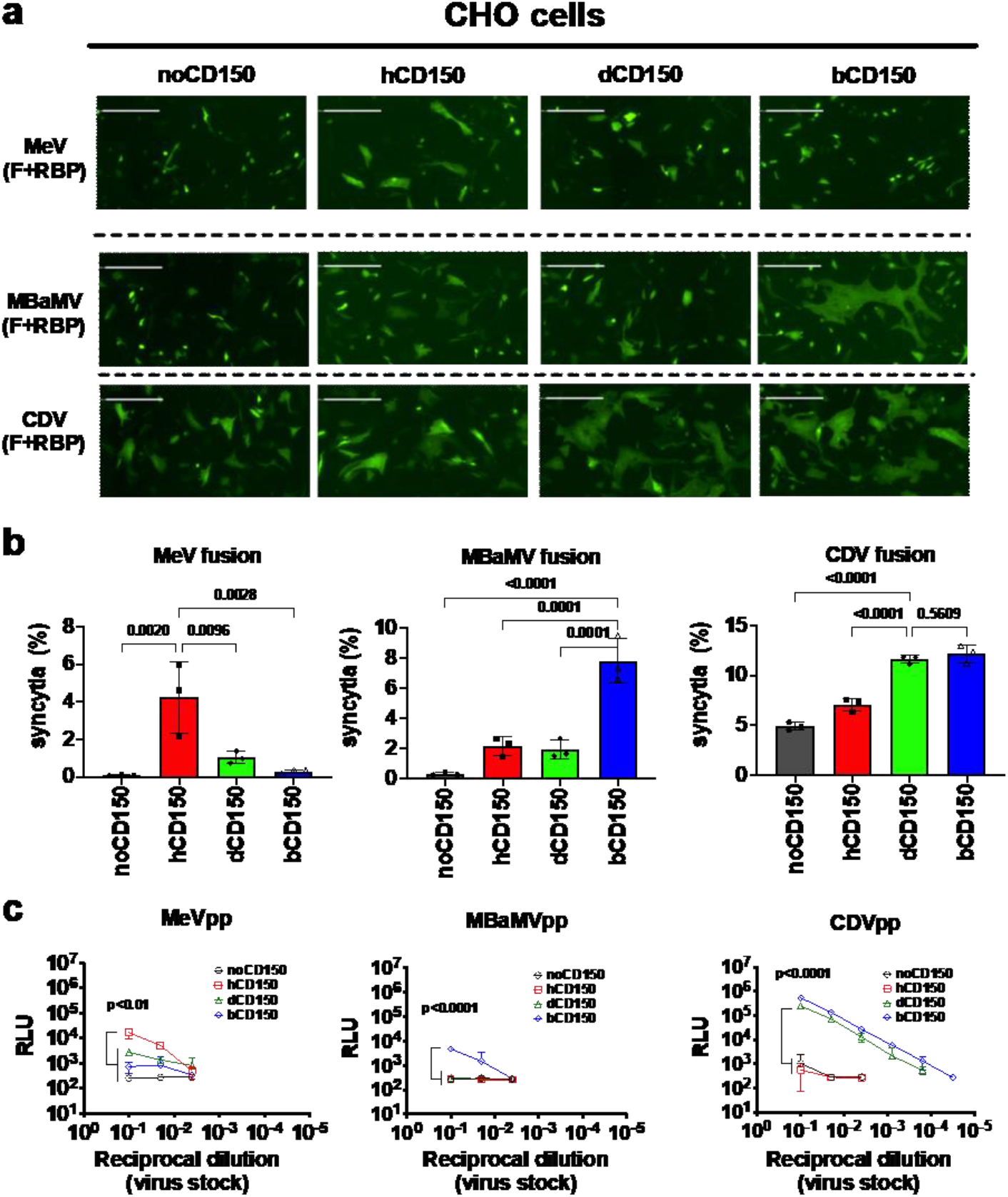
MBaMV envelope glycoproteins use host specific CD150 (SLAMF1) for fusion and entry. **a**, Syncytia formation in CHO cells co-transfected with the indicated morbillivirus envelope glycoproteins, species-specific CD150, and Life-act-GFP. Images were taken by the Celigo Imaging Cytometer (Nexcelom) at 48 hours post-transfection (hpt) and are computational composites from an identical number of fields in each well. White bar equals 200 μm. Brightness and contrast settings were identical. **b**, Quantification of syncytia formation in (a). Data are mean +/−S.D. from 3 independent experiments. Indicated adjusted p values are from ordinary one-way ANOVA with Dunnett’s multiple comparisons test. **c**, VSV-pseudo particle (pp) entry assay showed similar trends. Adjusted p values obtained as in **(b)** but only for comparing groups at the highest viral inoculum used (10^−1^ reciprocal dilution).

We also evaluated the receptor usage of MBaMV in a VSV-pseudotype entry assay. VSV-ΔG[Rluc] bearing MeV-RBP and F entered hCD150-transfected CHO cells better than dCD150-, bCD150-, or mock-transfected cells (Fig. 1c) as expected. MBaMV-pseudotypes entered only bCD150-transfected CHO cells. CDV-pseudotypes showed good entry into dCD150- and bCD150-transfected, but not hCD150-transfected CHO cells. These results are generally consistent with our fusion assay results and support the species specificity of morbilliviruses. CDV has a high propensity to cross species barriers and can cause disease in multiple carnivore families such as large felids (e.g. lions, tigers), hyaenids (e.g. spotted hyenas), ailurids (e.g. red pandas), ursids (e.g. black bears), procyonids (e.g. raccoons), mustelids (e.g. ferrets), viverrids (e.g. civets), and even non-carnivore species such as javalinas (peccaries) and rodents (Asian marmots)^20^. CDV has also been implicated in multiple outbreaks in non-human primates (various *Macaca* species)^21–23^. The ability of CDV to use bCD150 and dCD150 with equal efficiency suggests potential for efficient transmission from carnivores into some chiropteran species if other post-entry factors do not present additional restrictions.

### Generation of MBaMV by reverse genetics

Next, we attempted to generate a genomic cDNA clone of MBaMV that we could rescue by reverse genetics. We synthesized and assembled the putative MBaMV genome in increasingly larger fragments. Two silent mutations were introduced in the N-terminal 1.5 kb of the L gene to disrupt a cryptic open reading frame in the minus strand (Extended Data Fig. 3) that initially prevented cloning of the entire MBaMV genome. We introduced an additional EGFP transcription unit at the 3’ terminus and rescued this MBaMV-GFP genome using the N, P, and L accessory plasmid from MeV (Extended Data Fig. 1a). MBaMV-GFP was initially rescued in BSR-T7 cells but passaged, amplified, and titered on Vero-bCD150 cells (Extended Data Fig. 4a). MBaMV formed GFP-positive syncytia containing hundreds of nuclei at 3 days post-infection (dpi) (Fig. 2a) and relatively homogenous plaques by 7 dpi (Fig. 2b). Transmission electron microscopy (TEM) (Fig. 2c) captured numerous virions budding from Vero-bCD150 cells with pleiomorphic structure and size (~100-200 nm) consistent with paramyxovirus particles. At high magnification, virions were outlined by protrusions suggestive of surface glycoproteins. RNP-like structures can be found in the interior of the virion shown. These observations are consistent with previous findings from MeV^24^.

**Figure 2.**
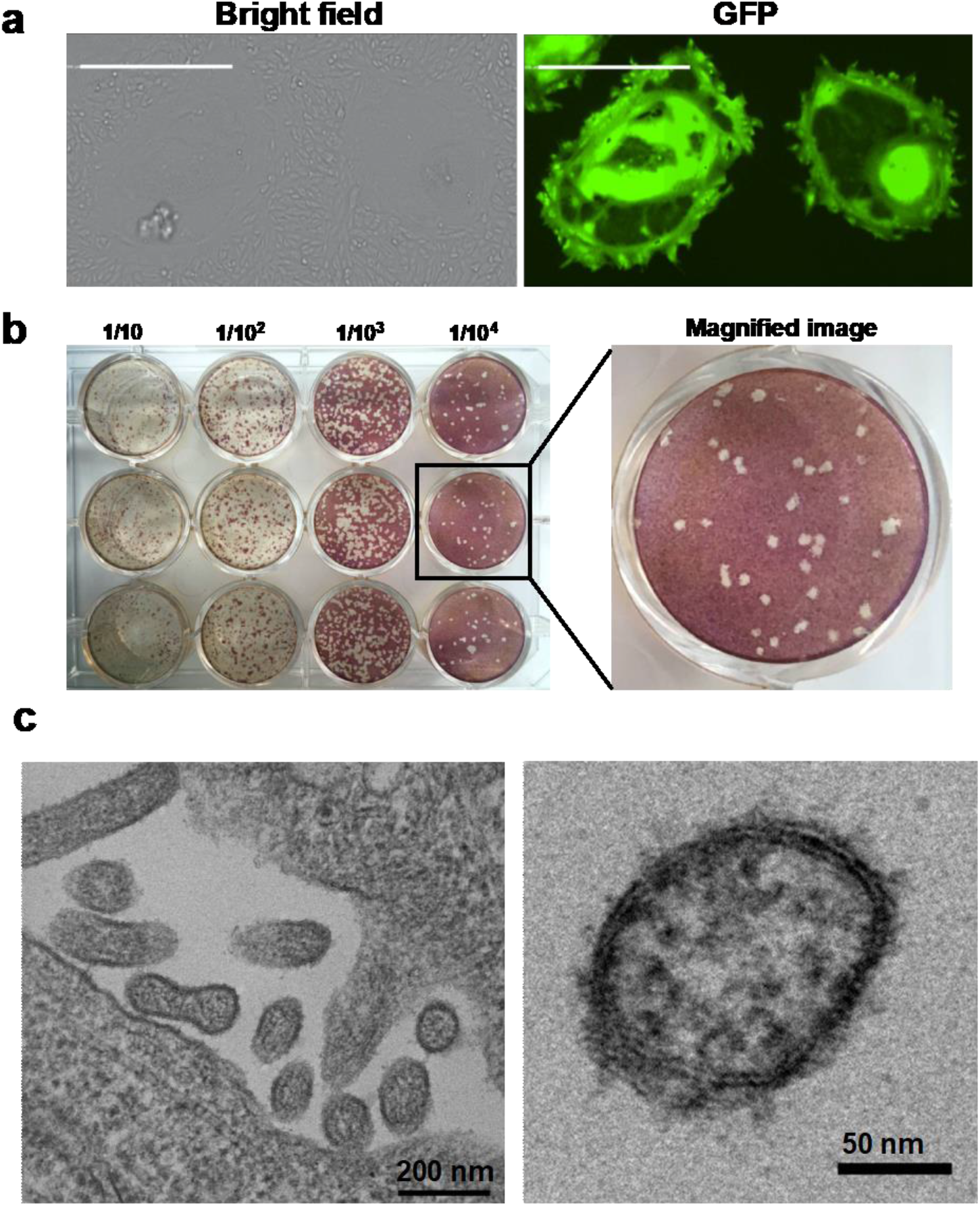
Virological characterization of myotis bat morbillivirus (MBaMV). **a**, Syncytia formation in Vero-bCD150 cells induced by MBaMV 3 days post-infection (dpi). Cells formed syncytia involving > 100 nuclei upon infection (bright field), which is clearly outlined by virus expressed GFP (right). Scale bar equals 500 micrometers. **b**, MBaMV plaque formation in Vero-bCD150 cells. Cells were infected by 10-fold serially diluted virus stock, incubated with methylcellulose containing-DMEM and stained with crystal violet and neutral red 7 dpi. Diameter of well is 22 mm. One well is magnified to show the plaque morphology in detail. **c**, shows transmission electron microscopy (TEM) images of MBaMV virion on the surface of Vero-bCD150 cells at 3 dpi. Numerous enveloped virions are budding from the plasma membrane (left). Magnified image (right) shows virion and ribonucleoprotein complex (RNP).

### Evaluation of receptor usage by MbaMV

To understand how well CD150 from various hosts supports MBaMV replication, we tested MBaMV growth in parental Vero-CCL81 cells and isogenic derivatives constitutively expressing CD150 of human, dog, or bat. MBaMV formed huge syncytia (Fig 3a) at 2 dpi in Vero-bCD150 cells and reached peak titers of ~10^5^ PFU/ml at 3 dpi (Fig 3b). MBaMV showed moderate syncytia spread and growth in Vero-dCD150 cells but peak titers at 5 dpi was ~100-fold lower. No significant virus growth was detected in Vero or Vero-hCD150 cells. These results confirm that MBaMV can use bCD150 but not hCD150 for efficient cell entry and replication. MBaMV appears to use dCD150, albeit to a much lesser extent than bCD150.

MeV uses human nectin-4 as the epithelial cell receptor^25,26^ which mediates efficient virus shedding from the affected host^16,27^. CDV also uses human nectin-4 efficiently for entry and growth^23^. To test if MBaMV can use human nectin-4 in an epithelial cell context, we evaluated the replication kinetics of MBaMV in human lung epithelial cells that express high (H441) or low (A549) levels of nectin-4^16,28^(Extended Data Fig. 4b). Surprisingly, MBaMV showed efficient virus spread (Fig. 3c) in H441 cells and reached 10^4^ PFU/ml by 6 dpi (Fig. 3d). In contrast, MBaMV showed small GFP foci and 10 times lower titer in A549 cells. Comparing the Area Under Curve (AUC) revealed significant differences in this growth curve metric (Fig. 3e). However, MeV still replicated to higher titers than MBaMV in H441 cells (Fig. 3d-e). This could be due to species specific host factors or differences in interferon antagonism between human and bat morbilliviruses. Thus, we tested MBaMV versus MeV growth in interferon-defective Vero-human nectin-4 cells (Vero-hN4). MBaMV and MeV replicated and spread equally well on Vero-hN4 cells (Fig 3f-g), validating the ability of MBaMV to use human nectin-4, and suggesting that MBaMV may not be able to counteract human innate immune responses.

**Figure 3.**
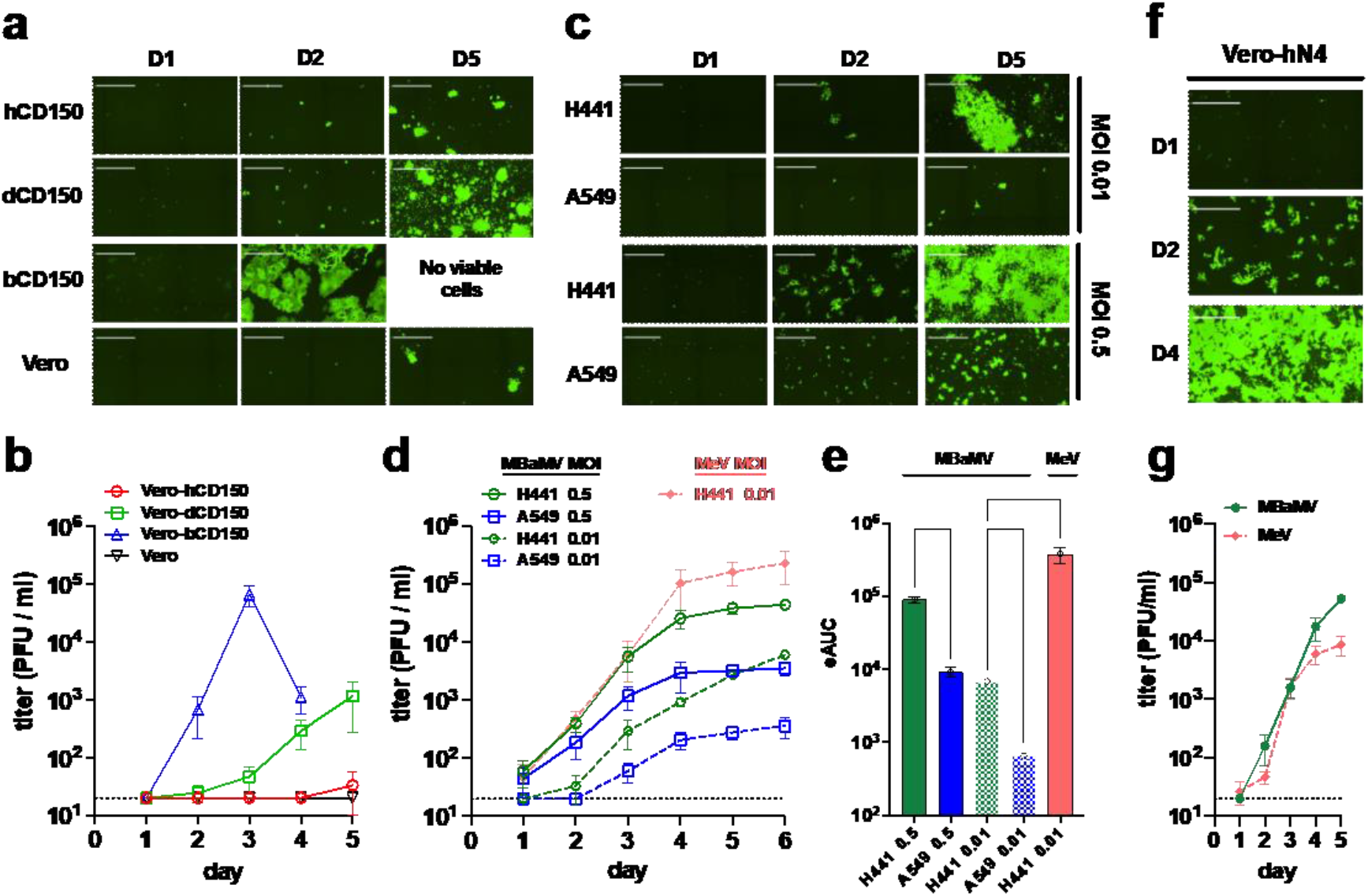
MBaMV replicates efficiently in cells expressing bCD150 and human nectin-4. **a-b**, Vero-hCD150, Vero-dCD150, Vero-bCD150, and Vero cells were infected with rMBaMV-EGFP (MOI 0.01). Virus replication and spread were monitored by imaging cytometry **(a)** and virus titer in the supernatant **(b). a**, Large syncytia were evident in Vero-bCD150 cells by 2 dpi. **b**, Supernatant was collected every day and the virus titer was determined by a GFP plaque assay (see methods). Data shown are mean +/−S.D. from triplicate experiments. **c-e**, H441 and A549 cells were infected with rMBaMV-EGFP at a low (0.01) or high (0.5) MOI. Virus replication and spread were monitored as in **a**-**b. c**, Infected H441 and A549 cells at 1, 2 and 5 dpi (D1, D2, D5). **d**, Virus growth curves represented by daily titers in the indicated conditions. Data shown are mean titers +/−S.D. from triplicate infections. **e**, The empirical Area Under Curve (eAUC) was obtained from each growth curve and plotted as a bar graph (mean +/−S.D.) (PRISM v 9.0). Adjusted p values are indicated (one-way ANOVA Dunnett’s T3 multiple comparison test). **f-g**, Vero-human nectin-4 cells (Vero-N4) were infected with MBaMV and MeV (MOI 0.01). **f**, MBaMV infected Vero-hN4 at D1, D2 and D4. **g**, Replicative virus titers for MBaMV and MeV on Vero-hN4 cells over 5 days (mean +/−S.D., n=3). White bar in **a, c**, and **f** equals 1 millimeter. All images shown are captured by a Celigo Imaging Cytometer (Nexcelom). Images are computational composites from an identical number of fields in each well. The limit of detection for virus titer determination is 20 PFU/ml and is indicated by the dotted line in **b, d, and g**.

### Molecular characterization of MbaMV

To better understand the transcriptional profile of MBaMV, we used Nanopore long-read direct RNA sequencing to sequence the mRNAs of MBaMV-infected Vero-bCD150 cells at 2 dpi (MOI=0.01). We found a characteristic 3’-5’ transcriptional gradient where GFP>N>P>M>F>RBP>L (Extended Data Fig. 5a). Morbilliviruses have a conserved intergenic motif (CUU) between the gene end and gene start of adjacent genes ‘AAAA-<u>CUU</u>-AGG’. This intergenic motif was not immediately apparent in the long complex M-F intergenic region of the assembled MBaMV genome. However, the high coverage of this M-F intergenic region (M read-through transcripts) identified the M-F intergenic motif as ‘CGU’ instead of ‘CUU’ (Extended Data Fig. 5b).

The P gene of morbilliviruses is known to generate the V or W genes through the insertion of one or two guanines, respectively, at the conserved editing motif (AAAAGGG)^29^, which is present in MBaMV. Amplicon sequencing of the P gene editing motif—from the same mRNA pool used above—revealed the frequency of P, V, and W mRNA is 42.1%, 51.2%, and 2.6%, respectively (Extended Data Fig. 5c), suggesting that the major interferon antagonist (V) is produced. This P-editing ratio is similar to what has been found in previous studies ^30^.

We next evaluated the expression and cleavage of two surface glycoproteins (RBP and F). C-terminal AU-1 tagged F construct showed uncleaved F0 and cleaved F1 (Extended Data Fig. 5d). C-terminal HA tagged RBP construct showed monomer in addition to oligomers (Extended Data Fig. 5e). MBaMV-RBP showed smear above 110 kDa which is suggestive of oligomerization. This oligomerization was also seen with MeV-RBP but not with CDV RBP suggesting differential stability under the sub-reducing conditions used.

### Species tropism of MbaMV

The two suborders of chiropterans (bats), Pteropodiformes (Yinpterochioptera) and Vespertilioniformes (Yangochiroptera), include more than 1,400 species grouped into 6 and 14 families, respectively^31^. Myotis bats belong to the prototypical Vespertilionidae family that is the namesake of its suborder. Jamaican fruit bats (*Artibeus jamaicensis*) belong to the same suborder as myotis bats, albeit from a different family (Phyllostomidae). We inoculated 6 Jamaican fruit bats available in a captive colony via two different routes with MBaMV to assess its pathogenicity *in vivo*. All bats remained asymptomatic and showed no evidence of developing systemic disease up to 3 weeks post-infection. Nor could we detect any molecular or serological evidence of productive infection (Extended Data Fig. 6). Inspection of Jamaican fruit bat and myotis CD150 sequences revealed key differences in the predicted contact surfaces with RBP (discussed below), which we speculate are responsible for the species-specific restriction seen in our experimental challenge of Jamaican fruit bats with MBaMV. To identify RBP-CD150 interactions likely involved in determining host species tropism, we compared the amino acid sequences at the putative contact surfaces of morbillivirus RBPs and their cognate CD150 receptors. Using PDBePISA^32^, we identified three key regions in MeV-RBP (residues 188-198, 498-507, and 524-556, Extended Data Fig. 7a-c) occluding two regions in CD150 (residues 60-92 and 119-131 of human CD150, Extended Data Fig. 8) in the crystal structure of MeV-RBP bound to CD150 (PDB ID: 3ALW)^33^. Alignment of key regions in morbillivirus RBPs implicated in CD150 interactions reveals virus-specific differences that suggest adaptation of morbillivirus RBPs to the CD150 receptors of their natural host. Most notably, MBaMV lacks the DxD motif at residues 501-503 (505-507 in MeV) that is present in all morbilliviruses except FeMV (Extended Data Fig. 7). These residues form multiple salt bridges and hydrogen bonds that stabilize MeV-RBP and hCD150 interactions. Their conservation suggest they perform similar roles for other morbilliviruses. On the CD150 side (Extended Data Fig. 8), residues 70-76 and 119-126 are the most variable between host species. Interestingly, Jamaican fruit bat and *Myotis* CD150 differ considerably in these regions, providing a rationale for the non-productive infection we saw in our Jamaican fruit bat challenge experiments.

### Susceptibility of human myeloid and lymphoid cells to MBaMV

Alveolar macrophages and activated T- and B-cells expressing CD150 are the initial targets for measles virus entry and systemic spread. To better assess the zoonotic potential of MBaMV, we compared how well human and bat morbilliviruses can infect human monocyte-derived macrophages (MDMs) and peripheral blood mononuclear cells (PBMCs). Both MeV and MBaMV infected MDMs were clearly GFP+ 24 hpi (Fig. 4a), but infection was variable between donors and even between different viral stocks on the same donor (Fig. 4b and 4d). However, MeV infection of MDMs was inhibited by sCD150 whereas MBaMV infection was not (Fig. 4c). MDMs had variable expression of CD150 (10-30% CD150+) but morbillivirus infection did not appear to be correlated with CD150 expression (Fig. 4d). Conversely, when PBMCs were stimulated with concanavalin A and IL-2, only MeV robustly infected these cells (Fig. 4e).

**Figure 4.**
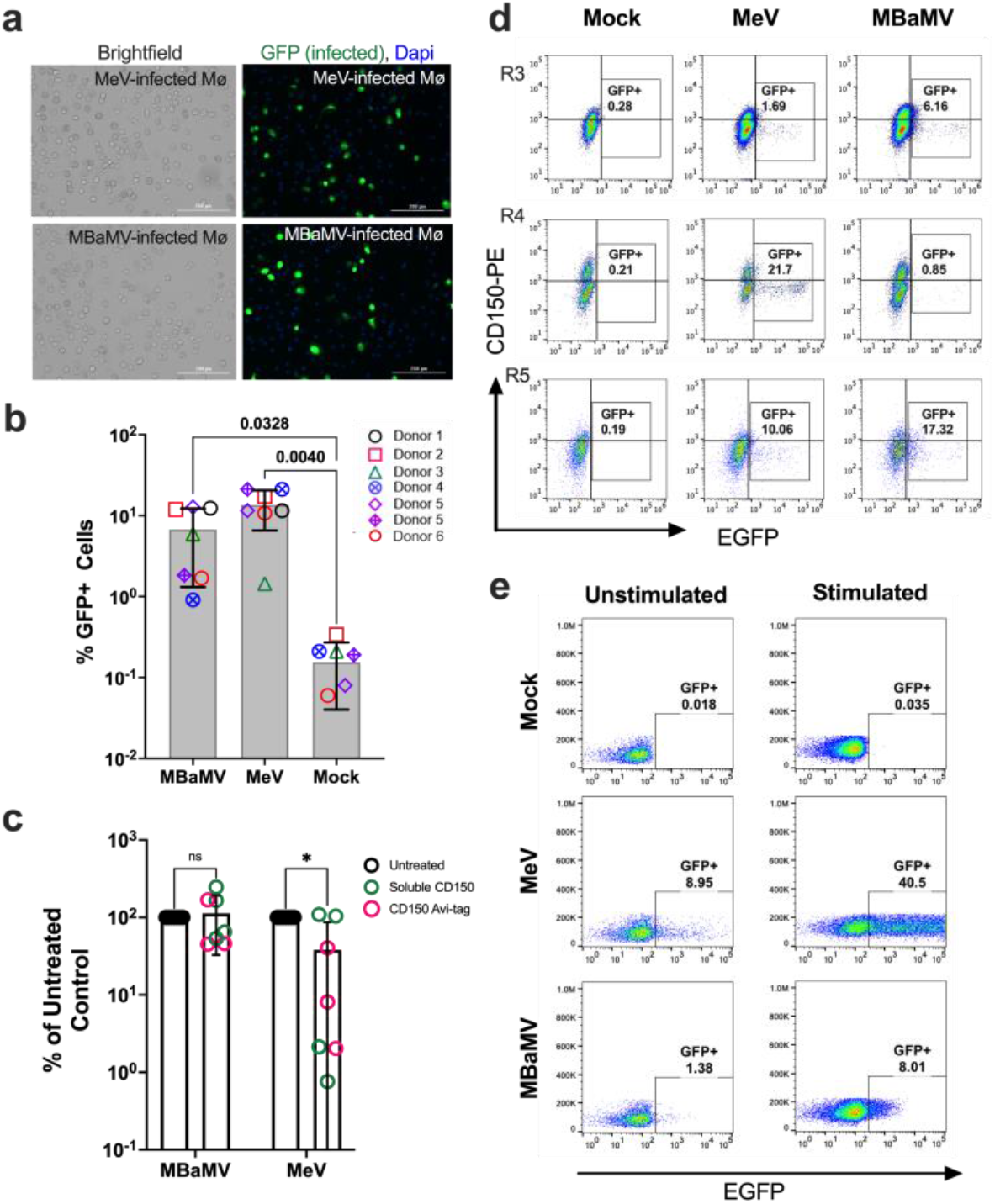
MBaMV infects human monocyte-derived macrophages (MDM) in a CD150-independent manner. **a-b**, MDMs were infected with MV323-EGFP or MBaMV (1×10^5^ IU/sample) and were either **(a)** fixed by 2% PFA at 24 hpi, DAPI-stained and imaged (scale bar is 200 μm), or **(b)** quantified by flow cytometry. The percent of CD68+GFP+ MDMs from 6 donors are shown. Open and crossed symbols indicate experiments using lot 1 and lot 2 viruses, respectively. Adjusted p values are from one way ANOVA with Dunnett’s multiple comparisons test. **c**, Soluble human CD150 (sCD150) or a CD150 Avi-tag inhibited MeV but not MBaMV infection of macrophages. GFP+ events in untreated controls were set to 100%, and entry under sCD150/ CD150 Avi-tag were normalized to untreated controls. Adjusted p values are from two-way ANOVA with Šídák’s multiple comparisons test. In (**b**) and (**c**), data shown are mean +/−S.D. from multiple experiments (N=5-7) with individual values also shown. **(d)** Exemplar FACS plots from the summary data shown in (**b**) for CD150 staining. **e**, ConA/IL-2 stimulated PBMCs were infected with MeV or MBaMV (MOI of 0.1) and analyzed for GFP expression by flow cytometry at 24 hpi.

### MBaMV infection may be blocked by human immune defenses

MeV-specific antibodies resulting from vaccination can provide cross protection against CDV infection^34^. To assess if human sera from MeV-vaccinated individuals could contain cross-neutralizing antibodies to MBaMV, we pooled MMR-reactive human sera and measured their ability to neutralize MeV, MBaMV, and CDV in hCD150, bCD150, and dCD150-expressing Vero cells. Human sera effectively neutralized MeV and MBaMV infection and, to a lesser extent, CDV infection (Fig. 5a, left and center panels). Conversely, sera from CDV-infected ferrets neutralized CDV infection much better than MeV or MBaMV (Fig. 5a, right panel). Sera from the MMR groups 1 and 2 had higher IC50s for MeV and MBaMV than for CDV while CDV-specific sera had a significantly higher IC50 for CDV than for MeV or MBaMV (Fig. 5b). These results indicate that human sera contain cross-neutralizing antibodies for MBaMV.

**Figure 5.**
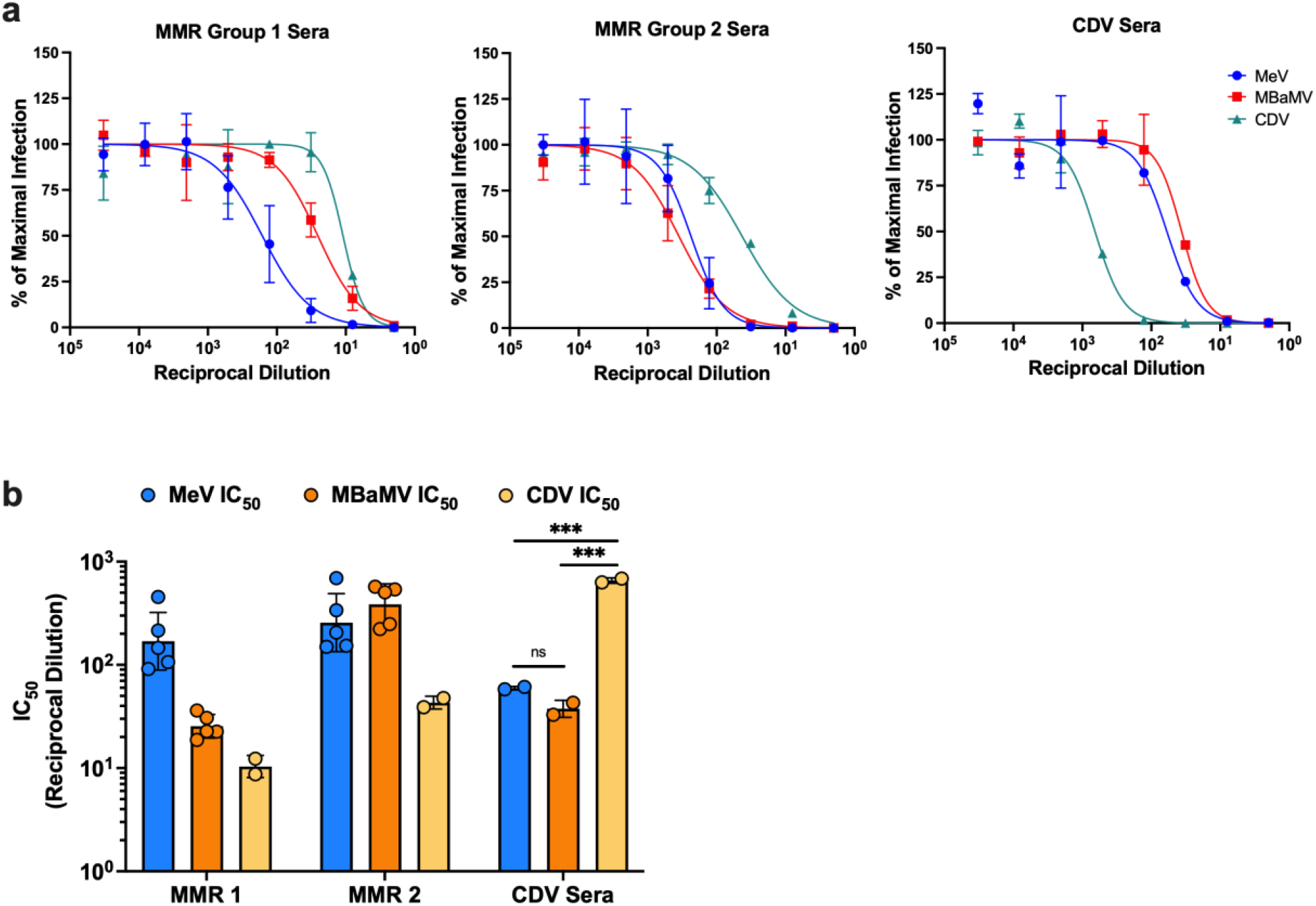
Human sera contains antibodies that partially cross-neutralize MBaMV. **a)** MeV, MBaMV, and CDV were incubated with serial dilutions of pooled human sera from MMR-vaccinated individuals (MMR group 1 and group 2) and sera from ferrets infected with CDV. The capacity for sera-treated virus to infect Vero cells expressing the appropriate receptor was measured by imaging infected cells 20 hpi, measuring the area of GFP+ cells, and calculating the reduction in infection compared to no sera controls. Neutralization curves were plotted for each virus and corresponding sera group. **b)** The IC50s from the neutralization curves shown in **a)** were generated for each replicate using a robust fit model and were plotted.

The MeV proteins P and V interfere with the innate immune system by disrupting the IFN pathway. Our sequencing results showed that MBaMV infection produced the P and V transcripts (Extended Data Fig. 5c), so we sought to determine if the MBaMV P and V proteins could antagonize the human IFN pathway. We found that cells transfected with MBaMV P or V and treated with IFN did not block ISRE induction, unlike ZIKV NS5, which effectively counteracts the ISRE (Extended Data Fig. 9a). Additionally, cells transfected with MBaMV P or V did not block the induction of IFN when treated with RIG-I, MDA5, or MAVS (Extended Data Fig. 9b-d). These data demonstrate that the MBaMV P and V proteins cannot antagonize the human IFN pathway.

### MBaMV is sensitive to morbillivirus RNA dependent RNA polymerase inhibitors

Potential drug treatments are a critical issue for emerging viruses. Thus, we tested if MBaMV is susceptible to currently available drugs. We have developed two orally bioavailable small compounds targeting the L protein of morbilliviruses, GHP-88309^35^ and ERDRP-0519^36^. The differences between MeV and MBaMV across the five functional domains of the L protein are shown schematically in Figure 6a^37^. *In silico* modelling (Fig. 6b) predicts that both drugs should bind similarly to MeV and MBaMV L protein. Closer inspection of the ERDRP-0519 binding pocket (Fig. 6c) shows 1155-1158 YGLE and H1288 residues interacting with ERDRP-0519.

These residues directly interact with ERDRP-0519 in MeV L^38^. Modeling of the GHP-88309 binding pocket (Fig. 6d) reveals involvement of E863, S869, Y942, I1009, and Y1105 residues which were previously reported as escape mutants of GHP-88309 in MeV^35^. As predicted, both drugs inhibited MBaMV growth in dose dependent manner (Fig. 6e and 6f). Although the EC_50_ of GHP-88309 is lower for MeV than MBaMV, (0.6 μM and 3.0 μM, respectively), GHP-88309 reaches a plasma concentration of >30 μM in animal models, indicating this drug could be an effective inhibitor of MBaMV *in vivo*.

**Figure 6.**
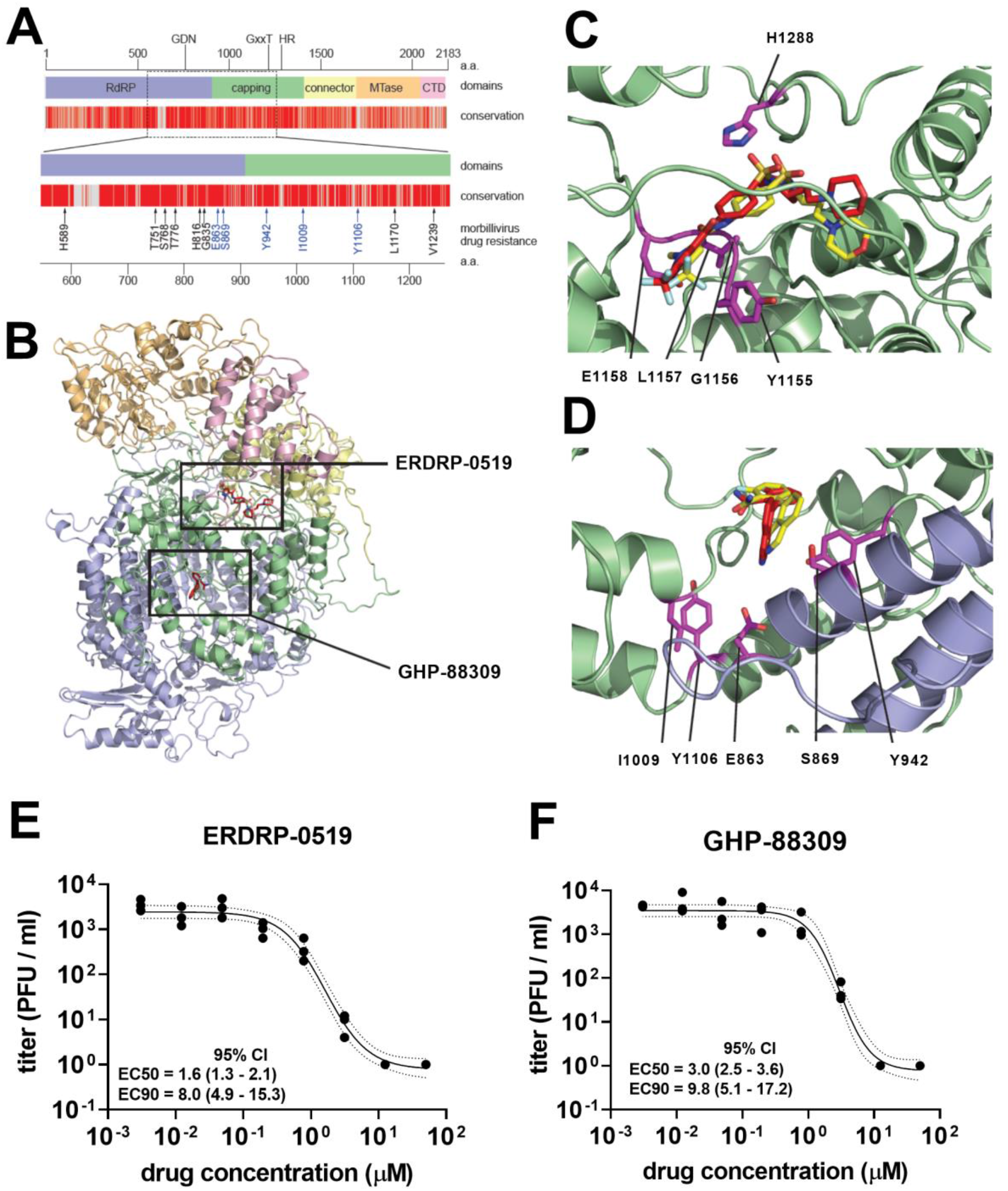
MBaMV is susceptible to RNA-dependent RNA polymerase inhibitor of ERDRP-0519 and GHP-88309. a, 2D-schematic of MBaMV L protein showing the layout of each domain. Conservation between the MeV and MBaMV L protein is shown. Differences between the MeV and MBaMV L proteins are shown as grey lines. b, A 3-D homology model of the MBaMV L protein was generated using the structural coordinates of the PIV5 L protein (PDB ID: 6V86). The RNA-dependent RNA polymerase (RdRP), capping, connector, methyltransferase (MTase), and C-terminal (CTD) domains are colored blue, green, yellow, orange, and pink, respectively. The locations of the top scoring in silico docking poses for ERDRP-0519 and GHP-88309 are boxed and the compounds are shown as red sticks. c, The top scoring docking pose of ERDRP-0519 in the homology model of MBaMV L protein (red sticks). An overlay of the previously identified docking pose of ERDRP-0519 in a homology model of MeV L protein is shown (yellow sticks) (Cox et al, PLoS Pathog, 2021. PMID:33621266). Residues identified in previous photoaffinity crosslinking experiments (Y1155, G1156, L1157 and E1158) and H1288 of the HR motif are shown as magenta sticks. d, The top scoring docking pose of GHP-88309 in the homology model of MBaMV L protein (red sticks). An overlay of the previously identified docking pose of GHP-88309 in a homology model of the MeV L protein is shown (yellow sticks) (Cox et al, Nature Microbiol, 2020, PMID:32661315). Residues identified in MeV resistance profiling studies are shown as magenta sticks. e, shows the dose-response inhibition growth curves of ERDRP-0519 against MBaMV. Vero-bCD150 cells were infected by MBaMV at MOI = 0.01 for 1 hour, then inoculum was replaced by fresh media containing inhibitor at the indicated concentrations (0 to 50 micromolar). 2 dpi, viral supernatants were collected and tittered on Vero-bCD150 cells as described in methods. Dots represent the values from 3 independent experiments. Regression curve (solid line) and 95% CI (dot line) were generated in PRISM (v.8.0). f, shows the drug response of GHP-88309 against MBaMV growth. The virus inhibition was conducted identically as for ERDRP0519.

## Discussion

Metagenomic viral surveillance studies aided by next-generation sequencing have allowed scientists to monitor viruses circulating in animal species and identify potential zoonotic threats^5– 7,39^. Surveillance of bat species has been particularly critical. For instance, >60 novel paramyxovirus sequences were identified in a 2012 bat surveillance study, several of which mapped to the *Morbillivirus* genus^4^. Recent metagenomic surveys confirm that bats harbor diverse orthoparamyxoviruses ^5–7^. While comparing novel virus sequences to known pathogens may help inform the risks associated with future spillover events, this type of *in silico* modeling based on viral sequences should also be complemented by functional characterization of such viruses. In this study, we identified a full-length morbillivirus genomic sequence from *Myotis riparius* bats in Brazil and generated an infectious virus clone using reverse genetics. With this approach, we circumvented the arduous process of isolating and culturing live virus directly from animals and instead produced MBaMV in the lab.

### MBaMV characterized as a morbillivirus

Prior to this study, there were only 7 ICTV recognized morbilliviruses species, none of which were isolated from bats. While the annotated MBaMV genome aligned with the classic morbillivirus genome organization (N, P/V/C, M, F, RBP, and L), it was important to verify that virus generated by reverse genetics successfully recapitulated morbillivirus biology. Fusion assays and entry experiments confirmed that MBaMV preferentially used myotis CD150 over human or dog CD150 to enter transgenic Vero cells (Fig. 3), which fits the paradigm that CD150 is the major determinant of host specificity for morbilliviruses. We also assessed P-editing—a hallmark of paramyxoviruses—and found RNA editing of P-mRNA, creating V-mRNA (single G insertion) or W-mRNA (double G insertion) of MBaMV. Interestingly, the proportion of V-mRNA at 51.2% of total P transcripts is unusually high for orthoparamyxoviruses, resembling the now extinct rinderpest virus (RPV) more than extant morbilliviruses^40^.

In their natural hosts, morbillivirus are highly pathogenic and can cause deadly acute infections^41^. Thus, one reasonable prediction is that MBaMV would cause visible disease in the bat host. However, when we challenged Jamaican fruit bats with MBaMV, we found the virus was *not* able to cause systemic disease in the bats (Extended Data Fig. 6) and there was no evidence that MBaMV productively infected these bats. This lack of infection could be due to the CD150 differences between the species—CD150 of Jamaican fruit bats and *Myotis* species is only 70% conserved on the amino acid level (Extended Fig. 8). We predict that MBaMV infection is more likely to cause serious disease in the *Myotis riparius* species. Alternatively, it is also possible that bat morbilliviruses do not cause severe illness in their hosts since bats possess unique immune systems that allow them to harbor deadly viruses such as Nipah, Ebola, and SARS without exhibiting illness.^42^

### Zoonotic potential of MBaMV?

When assessing the zoonotic potential of a novel virus, multiple factors must be considered, including receptor usage, the existence of cross-neutralizing antibodies in human sera, and interactions with the innate immune system. While non-human morbilliviruses are not currently known to jump the species barrier and infect humans, we did find that MBaMV was able to utilize human receptors *in vitro* to a certain extent. Traditionally, morbilliviruses use CD150 to enter myeloid and lymphoid cells. However, unlike MeV which infects human macrophages via CD150, MBaMV infects human macrophages in a CD150-independent manner (Fig. 4c)^43^. This result indicates that a non-CD150/nectin-4 entry receptor for MBaMV exists on human macrophages. In addition, MBaMV replicated well in H441 cells and in Vero cells expressing human nectin-4 (Fig. 3). CDV is also reported to use human nectin-4^23^ and can replicate in H358 cells^44^. Alarmingly, there have been several outbreaks of CDV in non-human primates, resulting in acute disease or death in the animals^34^. In one outbreak, mutations were found in the RBP which rendered CDV-RBP capable of efficiently using primate-CD150^23^. However, CDV is unlikely to adapt to humans in the presence of cross-reactive MeV immunity. Human sera from MMR-vaccinated individuals was able to cross-neutralize MBaMV infection *in vitro* (Fig. 5)—this would likely limit MBaMV infection if MMR-vaccinated humans were exposed to the virus. Finally, while MeV P and V proteins antagonize the innate immune response, MBaMV P and V were unable to block the IFN induction or signaling (Extended Data Fig. 9). Taken together, our findings suggest that the zoonotic potential for MBaMV is low due to cross-neutralizing anti-MeV antibodies and innate immune restriction. The former reinforces the need to maintain broad and high coverage of measles vaccination even when the virus has been eliminated in human populations.

In summary, our study provides a functional screening pipeline for evaluating the zoonotic potential of a paramyxovirus identified only from metagenomic data. Our comprehensive characterization will facilitate the screening of many other morbilliviruses present in bat and rodent reservoirs, including at least one other full-length morbillivirus sequence present in the greater spear-nosed bat (*Phyllostomus hastatus*). Given the deluge of metagenomic data from wild-life surveillance studies, a formal blueprint for evaluating the zoonotic potential of paramyxoviruses known to cause disease in humans is urgently needed.

## Materials and methods

### Method to isolate bat morbillivirus sequence

The bat surveillance was conducted in the Amazon region of Brazil. The bat was a subadult male (immature, but independent) and apparently healthy. Mitochondrial DNA profiling (MW554523 and MW557650) identified the bat as a riparian myotis (*Myotis riparius*). RNA was subjected to NGS analysis, and viral genome (MW557651) was assembled from fastq read files (GSE166170). The bat was captured by mist net, then oral, rectal, and urogenital swabs were all collected for RNA extraction. Total nucleic acid (TNA) was extracted using the Roche MagNA Pure 96 platform following the manufacturer’s protocol, then TNA was DNase treated (DNase I; Ambion, Life Technologies, Inc.) and reverse transcribed using SuperScript III (Invitrogen, Life Technologies, Inc.) with random hexamer primers. The cDNA was treated with RNase H before second-strand synthesis by Klenow fragment (3′ to 5′ exonuclease) (New England Biolabs), then the double-stranded cDNA was sheared into average of 200 bps fragments using a Covaris focused ultrasonicator E210. Sheared cDNA was deep sequenced using the Illumina HiSeq 2500 platform and reads were bioinformatically de novo assembled using MEGAHIT v1.2.8 after quality control steps and exclusion of host reads using Bowtie2 v2.3.5^45^. This method was same as previously published. The virus was identified in the rectal swab. MBaMV was identified as part of a metagenomic survey of bats sampled in Brazil and Malaysia. All the metadata associated with this study, including the number and species of bats sampled can be found in Wells et al.^7^

### Generation of phylogenetic tree and conservation matrix table

Amino acid sequences of L proteins were aligned by ClustalW, then the evolutionary history of L proteins was inferred by Maximum Likelihood method with bootstrap test of 1,000 replicates. All processes were done in MEGA X^46^. For conservation matrix table, amino acid sequences of each gene were aligned by ClustalW, then the conservations were evaluated. The accession numbers used for the alignment were summarized in Table S1.

### Cells

293T cells (ACTT Ca# CRL-3216), A549 cells (ATCC Ca# CCL-185), Vero cells (ATCC Cat# CCL-81, RRID:CVCL_0059), and BSR T7/5 cells (RRID:CVCL_RW96) were grown in in Dulbecco’s modified Eagle’s medium (DMEM, ThermoFisher Scientific, USA) supplemented with 10% fetal bovine serum (FBS, Atlanta Biologicals, USA) at 37°C. NCI-H441 cells (ATCC Ca# HTB-174) were grown in RPMI 1640 medium (ThermoFisher Scientific, USA) with 10% FBS. Vero-hCD150 (Vero-human SLAM) cells are Vero cells derivative which constitutively express hCD150. Vero-dCD150 cells are Vero cells derivative which constitutively express HA-dCD150. Vero-hCD150 cells^47^ and Vero-dCD150 cells^48^ were provided by Dr. Yanagi at Kyushu University and maintained in DMEM with 10% FBS. Vero-bCD150 cells and Vero-human nectin-4 cells were generated as written below and maintained in DMEM with 10% FBS. CHO cells were grown in DMEM/F12 (1:1) medium (gibco) with 10% FBS.

### Plasmids

We cloned the open reading frame of hCD150, dCD150, and bCD150 (from *Myostis brandtii* since the CD150 sequence from *M. riparius* is unknown) into the pCAGGS vector cut by *Eco*RI (NEB) and *Nhe*I-HF (NEB). We introduced HA tag-linker-Igk signal peptides (amino acids corresponding to; MVLQTQVFISLLLWISGAYG-YPYDVPDYA-GAQPARSP) at the N-terminus of CD150s as previously reported^49^. The sequence of hCD150, dCD150, bCD150 sequence were from NP_003028.1, NP_001003084.1, and XP_014402801.1, respectively. We synthesized codon optimized gene sequences at GeneArt Gene Synthesis (Invitrogen), generating pCAGGS-Igk-HA-hCD150, pCAGGS-Igk-HA-dCD150, pCAGGS-Igk-HA-bCD150. We also generated pCAGGS-Igk-HA-bCD150-P2A-Puro which additionally express puromycin resistant gene. For pCAGGS-human nectin-4-P2A-puro, synthesized DNA by GeneArt Gene Synthesis (Invitrogen) was cloned into pCAGGS.

The sequence of MBaMV RBP and F open reading frame were synthesized by GenScript. These were cloned into pCAGGS vector cut by *Eco*RI and *Nhe*I-HF with adding HA tag (RBP gene) or AU1 tag (F gene) in C-terminus, generating pCAGGS-MBaMV-RBP-HA, pCAGGS-MBaMV-F-AU1.

For MeV RBP and F expressing plasmid, we amplified RBP and F sequence from p(+) MV323-AcGFP with the addition of HA-tag and AU1-tag same as MBaMV-RBP and -F, creating pCAGGS-MeV-RBP-HA, pCAGGS-MeV-F-AU1. For CDV RBP and F cloning, we amplified RBP and F sequence from pCDV-5804P plasmid with the addition of HA-tag and AU1-tag, creating pCAGGS-CDV-RBP-HA, pCAGGS-CDV-F-AU1.

Genome coding plasmids for MeV; (p(+) MV323-AcGFP) and CDV; pCDV-5804P were kindly gifted from Dr. Makoto Takeda^50^ and Dr. Veronica von Messling respectively^51^. We transferred the MeV genome sequence into pEMC vector, adding an optimal T7 promotor, a hammer head ribozyme, and we introduced an eGFP transcriptional unit at the head of the genome (pEMC-IC323-eGFP), which is reported in the previous study^19^.

For the generation of MBaMV genome coding plasmid, we synthesized pieces of DNA at 2000 - 6000 bps at Genscript with the addition of eGFP transcriptional unit at the head of genome (eGFP-MBaMV). DNA fragments were assembled into pEMC vector one-by-one using in-fusion HD cloning kit (Takara), generating pEMC-eGFP-MBaMV. The N-terminal 1.5 kb of the L gene was initially unclonable. Sequence analysis revealed a putative 86 aa open reading frame (ORF-X) in the complementary strand. Introduction of two point mutations in this region to disrupt ORF-X without affecting the L amino acid sequence (Extended Data Fig. 4) finally enabled cloning of the full-length genome suggesting that ORF-X was likely toxic in bacteria.

### Recovery of recombinant MBaMV and MeV from cDNA

For the recovery of recombinant MBaMV, 4 × 10^5^ BSR-T7 cells were seeded in 6-well plates. The next day, the indicated amounts (written below) of antigenomic construct, helper plasmids (-N, -P and -L from measles virus), T7 construct, and LipofectamineLTX / PLUS reagent (Invitrogen) were combined in 200 mL Opti-MEM (Invitrogen). After incubation at room temperature for 30 minutes, the DNA - Lipofectamine mixture was added dropwise onto cells. The cells were incubated at 37°C for 24 hours. The cells containing P0 viruses were trypsinized and passed onto Vero-bCD150 cells (2.0×10^6^ cells / flask in one 75cm^2^ flask). We collected supernatant 2 days after overlay (P1 virus) and reamplified MBaMV in fresh Vero-bCD150 cells (> 2X T175 cm^2^ flasks). These passage 2 (P2) stocks were titerred, frozen down in aliquots, and used for all experiments. The amount of measles plasmids used for rescue is reported in our previous study^52^: 5 mg antigenomic construct, 1.2 mg T7-MeV-N, 1.2 mg T7-MeV-P, 0.4 mg T7-MeV-L, 3 mg of a plasmid encoding a codon-optimized T7 polymerase, 5.8 mL PLUS reagent, and 9.3 mL Lipofectamine LTX.

The rescue of MeV was done exactly same way as MBaMV rescue except that 5 mg of pEMC-IC323eGFP was used for transfection and Vero-hCD150 cells were used for coculturing.

### Titration of viruses and plaque assay

For MBaMV, a monolayer of Vero-bCD150 cells in 12 well was infected by 500 ml of serially diluted samples for 1 hour, followed by medium replacement with methylcellulose containing DMEM. 5 dpi, the number of GFP positive plaque was counted to determine titer. For the plaque assay, infected Vero-bCD150 cells were incubated under methylcellulose containing DMEM for 7 days. Cells were then stained with 1% crystal violet and 1% neutral red sequentially. For MeV, we used Vero-hCD150 cells and fixed the plates at 4dpi.

### Growth analysis

2.0 × 10^5^ cells / well were seeded in 12 well plate. Cells were infected by indicated titer of viruses (MOI 0.01 or 0.5) for one hour, followed by replacement of fresh medium. Viruses were grown for 5 days with medium change every day. Collected supernatants were used for titration.

### Generation of Vero-bCD150 cells and Vero-human nectin-4 cells

4.0 × 10^5^ of VeroCCL81 cells were transfected with 2 mg of pCAGGS-Igk-HA-bCD150-P2A-Puro with Lipofectamine 2000 (Invitrogen); cells were selected under 5 mg/ml of puromycin (Gibco) until colonies were visible. Colonies were isolated independently and checked for HA expression using FACS. Vero-human nectin-4 cells were generated by transfecting pCAGGS-human nectin-4-P2A-Puro into VeroCCL81 cells, followed by 5 mg/ml of puromycin selection, and clone isolation. Surface expression was checked by FACS.

### Generation of VSV-pseudotyped virus and entry assay

6 × 10^6^ cells of 293T were seeded in a 10cm dish (pre-coated by poly-L-lysine (Sigma)) one day before transfection. 12 mg of RBP plus 12 mg of F coding plasmid from MeV, CDV, or MBaMV were transfected to cells by PEI MAX (polysciences). Vesicular stomatitis virus (VSV)-deltaG-Gluc supplemented by G protein (VSVDG-G*) were infected at MOI = 10 for one hour at 8 hours post plasmid transfection. Cells were washed with PBS three times and medium was maintained with Opti-MEM for 48 hours. Supernatant was collected and ultra-centrifuged at 25,000 rpm x 2 hours and the pellet was re-suspended with 100ul of PBS^53^. For the quantification of pseudotyped viral entry, CHO cells in 10cm dish were transfected with 24 mg of hCD150, dCD150, or bCD150 expressing plasmid with PEI MAX. CHO cells were passaged onto 96 well plates at 8 hours post transfection The pseudotyped-VSV of MeV, CDV, or MBaMV were used to infect the CHO cells. *Renilla* luciferase units (RLU) were measured by *Renilla* luciferase assay system (Promega) to quantify the pseudotype virus entry into cells.

### Image based fusion assay

CHO cells were seeded at 50,000 cells in 48-well dish 24 hours before transfection. Cells were transfected with 200 mg of pCAGSS-RBP-HA (of MeV/CDV/MBaMV), 200 mg of pCAGGS-F-AU1(of MeV/CDV/MBaMV), pCAGGS-Igk-HA-CD150 (20 ng human, 5 ng dog, or 20 ng bat), and 50 mg of pEGFP-C1 Lifeact-EGFP (purchased from Addgene) with 2.5 ml of polyethylenimine max (polysciences). At 36 hours post transfection, cells were imaged with a Celigo imaging cytometer (Nexcelom) with the GFP channel, and pictures were exported at the resolution of 5 micrometer / pixel. The GFP-positive foci (single cell or syncytia) were analyzed by ImageJ (developed by NIH), creating the profile of individual GFP-positive foci with size information.

For the evaluation of syncytia size, we first filtered the GFP-positive foci with the size of >= 10 pixel^2^, which is the median size of GFP area in the well of MeV-F plus LifeactGFP transfection to exclude non-specific background noise. Then we calculated the frequency of syncytia which is defined as the GFP counts of >= 100 pixel^2^ (10 times of median size of single cells) / total GFP counts of >= 10 pixel^2^.

### Surface expression check of bCD150 in Vero-bCD150 cells and human nectin-4 in Vero-human nectin-4 cells by FACS

50,000 cells in a 96 well plate were dissociated with 10 μM EDTA in DPBS, followed by a 2% FBS in DPBS block. Cells were treated with primary antibody for one hour at 4°C, then washed and treated by secondary antibody for one hour at 4°C. Vero-bCD150 cells were examined with a Guava® easyCyte™ Flow Cytometers (Luminex) for the detection of signal. Vero-human nectin-4 cells were subjected to Attune NxT Flow Cytometer (ThermoFisher Scientific). For primary antibody, mouse monoclonal nectin-4 antibody (clone N4.61, Millipore Sigma) and rabbit polyclonal HA tag antibody (Novus biologicals) were used at appropriate concentration indicated by the vendors. For secondary antibody, goat anti-rabbit IgG H&L Alexa Fluor® 647 (Abcam) and goat anti-mouse IgG H&L Alexa Fluor® 647 (Abcam) were used appropriately. FlowJo was used for analyzing FACS data and presentation.

### Soluble CD150 production and purification

Production and purification of soluble CD150 is as previously reported^54^. Soluble CD150 is a chimera comprising the human V (T25 to Y138) and mouse C2 domains (E140 to E239) + His6-tag, which was cloned into pCA7 vector. The expression plasmid was transfected by using polyethyleneimine, together with the plasmid encoding the SV40 large T antigen, into 90% confluent HEK293S cells lacking N-acetylglucosaminyltransferase I (GnTI) activity. The cells were cultured in DMEM (MP Biomedicals), supplemented with 10% FCS (Invitrogen), l-glutamine, and nonessential amino acids (GIBCO). The concentration of FCS was lowered to 2% after transfection. The His6-tagged protein was purified at 4 days post transfection from the culture media by using the Ni2+-NTA affinity column and superdex 200 GL 10/300 gel filtration chromatography (Amersham Biosciences). The pH of all buffers were adjusted to 8.0. Soluble CD150 Fc fusion avitag was purchased from BPSbioscience, and reconstituted by PBS.

### Macrophage experiments

CD14+ monocytes were isolated from leukopaks purchased from the New York Blood Bank using the EasySep Human CD14 positive selection kit (StemCell #17858). For macrophage differentiation, CD14+ monocytes were seeded at 10^6^ cells/ml and cultured in R10 media (RPMI supplemented with FBS, HEPES, L-glutamine, and pen/strep) with 50 ng/ml of GM-CSF (Sigma Aldrich G5035) in a 37°C incubator. Media and cytokines were replaced 3 days post seeding. At 6 days post seeding, macrophages were infected with either MeV or MBaMV at 100,000 IU (infectious units) per 500,000 cells and were spinoculated at 1,200 rpm for 1 hour at room temperature. Virus inoculum was removed and cells were incubated in R10 media with GM-CSF at 37°C. For imaging experiments, macrophages were fixed in 4% PFA at 30 hours post infection (hpi), stained with DAPI, and fluorescent and bright field images were captured on the Cytation 3 plate reader. For flow cytometry experiments, infected macrophages were stained for viability at 24 hpi (LIVE/DEAD fixable stain kit from Invitrogen L34976), treated with human Fc block (BD Biosciences), stained with antibodies against CD14 (eBioscience clone 61d3) and HLA-DR (eBioscience clone LN3), fixed in 2% PFA, permeabilized with saponin, and stained for intracellular CD68 (eBioscience clone Y1/82A) and CD150 (eBioscience). Soluble CD150 (1mg/ml) or CD150 Avi-tag (BPS Bioscience) were incubated with MeV or MBaMV for 15 minutes prior to infection for inhibition experiments. Stained macrophages were run through an Attune NxT Flow Cytometer and data was analyzed using FlowJo software (v10).

### T cell experiments

PBMCs were isolated from fresh blood donations obtained through the New York Blood Center using density centrifugation and a ficoll gradient. Isolated PBMCs were then resuspended in RPMI media (10% FBS, 1% L-Glutamine, 1% Penicillin-Streptomycin) and were stimulated for T-cell activation with Concanavalin-A (ConA) at 5 ug/ml for 72 hours. Following, cells were washed once with PBS and stimulated with 10 ng/ml of IL2 for 48 hours. Cells were subsequently infected at an MOI of 0.2 with MeV, BaMV or were mock infected in 12 well plates at 10^6^ cells/ml. Cells were collected 24 hours post infection, stained with Invitrogen’s LIVE/DEAD Fixable dead cell far red dye as per the manufacturer’s protocol, and were analyzed for eGFP expression by flow cytometry with an Attune NxT Flow Cytometer. Analysis was completed using FCSExpress-7. A total of 2 donors were utilized for this analysis, with the data from donor 1 shown in Figure 4.

### Western blot for RBP and F protein

1 × 10^6^ of 293T cells were seeded on to collagen coated 6 well plate. 293T cells were transfected by 2 mg of pCAGGS, pCAGGS-MBaMV-RBP-HA, or pCAGGS-MBaMV-F-AU1 using polyethylenimine max (polysciences). Cells were washed with PBS, then lysed by RIPA buffer. Collected cytosolic proteins were run on 4 - 15% poly polyacrylamide gel (Bio-rad. #4561086) and transferred onto PVDF membrane (FisherScientific, #45-004-113), followed by primary antibody reaction and secondary antibody reaction. Rabbit polyclonal HA tag antibody (Novus biologicals, #NB600-363), rabbit polyclonal AU1 epitope antibody (Novus biologicals, #NB600-453) was used for primary antibody for HA and AU1 tag detection. Rabbit monoclonal antibody (Cell signaling technology, #2118) were chosen as primary antibody to detect GAPDH. Alexa Fluor 647-conjugated anti-rabbit antibody (Invitrogen, #A-21245) was used as secondary antibody appropriately. Image capturing were done by Chemidoc™ MP (Biorad).

### Transcriptome analysis of MBaMV

4.0×10^5^ Vero-bCD150 cells were infected by MBaMV at MOI = 0.01. Cytosolic RNA was collected by 500 ml of Trizol (Ambion) at 2 dpi. Collected cytosolic RNA was sequenced by direct RNA sequence by MinION (Oxford Nanopore Technologies) with some modifications in the protocol. First, we started library preparation from 3 mg of RNA. Second, we used SuperScript IV (Invitrogen) instead of SuperScript III. Sequencing was run for 48 hours by using R9.4 flow cells. The fastq file was aligned to MBaMV genome sequence by minimap2 and coverage information was extracted by IGVtools.

### Evaluation of P mRNA editing

Infection and RNA extraction was same as above (transcriptome analysis). 1 ug RNA was reverse transcribed by TetroRT (bioline) with poly-A primer, followed by PCR with primer set of Pedit-f (sequence; GGGACCTGTTGCCCGTTTTA) and Pedit-r (sequence; TGTCGGACCTCTTACTACTAGACT). Amplicons were processed by using NEBNext Ultra DNA Library Prep kit following the manufacturer’s recommendations (Illumina, San Diego, CA, USA), and sequenced by Illumina MiSeq on a 2×250 paired-end configuration at GENEWIZ, Inc (South Plainfield, NJ, USA). Base calling was conducted by the Illumina Control Software (HCS) on the Illumina instrument. The paired-end fastq files were merged by BBTools. These merged fastq files were aligned to the reference sequence using bowtie2, creating a SAM file, and we counted the number of P-editing inserts.

### Neutralization Assay

Vero-hCD150, Vero-dCD150, and Vero bCD150 cells were seeded in 96-well plates. Two groups of pooled human sera from people who previously received the MMR vaccine (3 individuals per pool) and sera from ferrets infected with CDV (courtesy of Richard Plemper) were heat-inactivated for 30 minutes at 56°C. Equal amounts of CDV, MeV, and MBaMV (20,000 IU/mL) were incubated with serial dilutions of the heat-inactivated sera for 15 minutes at room temperature. Virus and sera were then added to the Vero cells with the correct receptor and placed at 37°C. At 20 hours post infection, the cells were imaged using a Celigo imaging cytometer (Nexelcom) with the GFP channel. Exported images were analyzed using ImageJ to measure the extent of viral infection by GFP+ area (MeV and MBaMV), or total GPF + counts (CDV). The % reduction in infection was calculated by setting the level of infection in the no sera control wells to 100%. The normalized data was plotted using GraphPad Prism and neutralization curves were generated using non-linear regression with [inhibitor] vs. normalized response. IC50 values were calculated for each replicate using a robust fit model. Five replicates were completed for the MeV and MBaMV neutralization with the pooled human sera and 2 replicates were repeated with CDV and the ferret sera.

### Interferon Induction and Response Assays

For ISG induction assays, HEK 293T cells were transfected with plasmids coding for ISG54-ISRE-FLuc, TK-RLuc, and either empty vector, MBaMV P, MVaMV V, or ZIKV MR766 NS5. At 24 hours post-transfection, the cells with treated with 100U of human IFNb (at 100U/mL). Cells were lysed 24 hours after IFNb treatment and FLuc and RLuc expression was measured using the Promega Dual luciferase assay. Data was calculated as a ratio of Fluc:RLuc to normalize for transfection efficiency. Two independent experiments with 3 technical replicates were completed. To measure the antagonism of IFN induction (IFNb promoter activation), HEK 293T cells were transfected with plasmids coding for IFNb-FLuc, TK-RLuc, an IFN promoter stimulant (either RIG-I, MDA5, or MAVS), and empty vector, MBaMV P, MBaMV V, or HCV NS3/4A (potential IFN antagonists). At 24 hours post transfection, cells were lysed and FLuc and RLuc expression was measured by Promega Dual luciferase assay. Data was calculated as a ratio of Fluc:RLuc to normalize for transfection efficiency. Two independent experiments with 3 technical replicates were completed. For statistical analysis, one-way ANOVA with Dunnett’s multiple comparisons were performed with Prism.

### Bat challenge experiment and evaluation of infection

Six Jamaican fruit bats (*Artibeus jamaicensis*) were inoculated with 2×10^5^ PFU MBaMV-eGFP; three bats were intranasally (I.N.) and 3 bats were intraperitoneally (I.P.). At 1 week post virus inoculation, bats were subjected to blood and serum collection, visually inspected for GFP expression around the nares, oral cavity, and eyes by LED camera in each group (I.N. and I.P.). At 2 weeks post virus infection, blood, serum, and tissues (lung, spleen, and liver) were collected from one bat in each group. At 3 weeks post virus infection, blood, serum, and tissues (lung, spleen, and liver) were collected from one bat in each group.

Blood RNA was extracted by Trizol. RNA was reverse transcribed by Tetro cDNA synthesis kit (Bioline) with the primer of ‘GAGCAAAGACCCCAACGAGA’ targeting MBaMV-GFP genome, then the number of genomes was quantified by SensiFAST™ SYBR® & Fluorescein Kit (Bioline) and CFX96 Touch Real-Time PCR Detection System (Biorad). The primer set for qPCR is ‘GGGGTGCTATCAGAGGCATC’ and ‘TAGGACCCTTGGTACCGGAG’.

Virus neutralization assay was done as follows. Heat inactivated (56°C x 30 minutes) bat serum was serially diluted by 3 times (starting from 5 times dilution) and mixed with 2 × 10^4^ PFU /ml of MBaMV at 1: 1 ratio for 10 minutes at room temperature. 100 ml of mixture was applied to Vero-batCD150 cells in 96 well. GFP foci were detected and counted by Celigo imaging cytometer (Nexcelom). GFP counts of serum treated samples were normalized by no serum treated well. Tissues were fixed with 10% buffered formalin and embedded with paraffin, then thin-sliced. GFP-IHC was performed by using VENTANA DISCOVERY ULTRA. Rabbit monoclonal antibody (Cell signaling technology, #2956) was used as a primary antibody, and OMNIMap anti-rabbit-HRP (Roche, #760-4310) was used as a secondary antibody. The GFP signal was visualized by using Discovery ChromoMap DAB kit (Roche, #760-2513). Tissues were counterstained with hematoxylin to visualize the nuclei.

### In-silico docking

In silico docking was performed with MOE 2018.1001 (Chemical Computing Group), as previously described^38^. A homology model of MBaMV L was created based on the structural coordinates of PIV5-L (PDB ID: 6V86) using the SWISS-MODEL homology modeling server^55^. Prior to docking, the model of the MBaMV L protein was protonated and energy minimized. An induced-fit protocol using the Amber10 force field was implemented to dock ERDRP-0519 and GHP-88309 into MBaMV L. For binding of ERDRP-0519, residues Y1155, G1156, L1157, E1158, and H1288 and for binding of GHP-88309, residues E858, D863, D997, I1009, and Y1106 were pre-selected as docking targets, which are predicted to line the docking sites of ERDRP-0519 and GHP-88309, respectively, in MeV L. Top scoring docking poses were selected and aligned in Pymol to the previously characterized in silico docking poses of the inhibitors to MeV L protein. Sequence alignment of MBaMV and MeV L proteins was performed using Clustal Omega^56^. Conservation was scored using the AL2CO alignment conservation server^57^.

### Transmission electron microscopy (TEM)

Routine transmission electron microscopy processing was done as described. The Vero-bCD150 cells infected by MBaMV for 3 days were washed with phosphate-buffered saline and then fixed with 2.5% glutaraldehyde in 0.1 M sodium cacodylate buffer (pH 7.4) on ice for 1 hour. The cells were scraped off the 100 mm tissue culture treated petri dish and pelleted by low-speed centrifugation (400g for 5 minutes). The pellet was fixed for 30 minutes with the same fixative before secondary fixation with 2% osmium tetraoxide on ice for 1 hour. The cells were then stained with 2% uranyl aqueous solution *en bloc* for 1 hour at room temperature, dehydrated with a series of increasing ethanol gradients followed by propylene oxide treatment, and embedded in Embed 812 Resin mixture (Electron Microscopy Sciences). Blocks were cured for 48 h at 65°C and then trimmed into 70 nm ultrathin sections using a diamond knife on a Leica Ultracut 6 and transferred onto 200 mesh copper grids. Sections were counterstained with 2% uranyl acetate in 70% ethanol for 3 min at room temperature and in lead citrate for 3 minutes at room temperature, and then examined with a JEOL JSM 1400 transmission electron microscope equipped with two CCD camera for digital image acquisition: Veleta 2K x 2K and Quemesa 11 megapixel (EMSIS, Germany) operated at 100 kV.

### Ethics declaration

Animal study was performed following the Guide for the Care and Use of Laboratory Animals. Animal experiment was approved by the Institutional Animal Care and Use Committee of Colorado State University (protocol number 1090) in advance and conducted in compliance with the Association for the Assessment and Accreditation of Laboratory Animal Care guidelines, National Institutes of Health regulations, Colorado State University policy, and local, state and federal laws. Archival CDV hyperimmune ferret sera were obtained from previous animal experiments approved by the Institutional Animal Care and Use Committee of Georgia State University (protocol number XXXX).

### Human subjects research

Normal primary dendritic cells and macrophages used in this project were sourced from ‘human peripheral blood Leukopack, fresh’ which is provided by the commercial provider New York Blood center, inc. Leukapheresis was performed on normal donors using Institutional Review Board (IRB)-approved consent forms and protocols by the vendor. The vendor holds the donor consents and the legal authorization that should give permission for all research use. The vendor is not involved in the study design and has no role in this project. Samples were deidentified by the vender and provided to us. To protect the privacy of donors, the vendor doesn’t disclose any donor records. If used for research purposes only, the donor consent applies. Aliquots of pooled immune sera were obtained from a previous anonymous serosurvey study that was qualified as Exemption 4 under NIH Exempt Human Subjects Research guidelines (Icahn School of Medicine at Mount Sinai).

### Data and materials availability

The raw next generation sequencing results of bat surveillance, P gene editing, and transcriptome by MinION are uploaded at NCBI GEO: GSE166170, GSE166158, and GSE166172, respectively.

Assembled MBaMV sequence and pEMC-MBaMVeGFP sequence information are available at MW557651 and MW553715, respectively. Cytochrome oxidase I host sequence and cytochrome b host sequence of virus infected bat are available at MW554523 and MW557650. MeV genomic cDNA coding plasmid (pEMC-IC323eGFP) sequence is available at NCBI Genbank: MW401770.

## Authors contributions

SI, SJA and BL conceived this study. SI conducted fusion assay, rescuing viruses, growth analysis, RNA sequencing of transcriptome analysis, and generation of cell lines written in the study. RLF conducted TEM imaging. JCC, JA, AP, and JL performed the macrophage and T cell experiments and data analysis. KYO conducted VSV-pseudotype entry assay. RMC and PKP provided ERDRP-0519 and GHP-88309 in addition to *in silico* modelling of MBaMV-L. HPC evaluated protein production by Western blot. TH provided structure-guided insights into conservation of RBP and CD150 binding as well as soluble human CD150 for inhibition assay. KYO and SK evaluated surface expression of morbillivirus receptors. CSS evaluated P-mRNA editing frequency from NGS data. TS, ME, SZ performed bat challenge experiment. ED conducted bat surveillance in collaboration with JEE and PD. SJA and HW conducted NGS analysis of bat surveillance and retrieved MBaMV sequences. ME and EV performed the IFN response and Induction experiments. JEE, PD and SJA provided insights into viral ecology and zoonotic threats. BL supervised this study. SI, JCC, SJA, and BL wrote the manuscript.

## Acknowledgements

S.I. was supported by Fukuoka University’s Clinical Hematology and Oncology Study Group (CHOT-SG). This study was supported in part by NIH grants AI123449 (B.L.), AI071002 (R.K.P and B.L.), AI149033 (B.L. and J.L.), AI140442 (T.S.), AI134768 (T.S.), and USAID PREDICT (S.J.A, J.H.E., P.D., E.D.). J.C.C. J.A.A., K.Y.O, A.P., C.S.S. acknowledge support from T32 AI07647. K.Y.O. was additionally supported by F31 (AI154739). J.A.A was additionally supported by F31 (HL149295). J.C.C. was additionally supported by F32 (HL158173). This work was also supported by Japan Agent for Medical Research and Development (AMED) Grant 20wm0325002h (T.H.), JSPS KAKENHI Grant Numbers 20H03497 (T.H.) and Joint Usage/Research Center program of Institute for Frontier Life and Medical Sciences, Kyoto University (S.I.).

