## Extended Data for "Assessing the zoonotic potential of a novel bat morbillivirus"

**a**

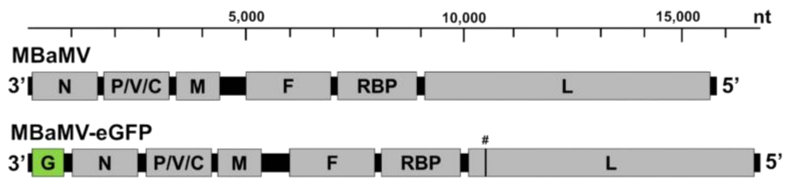


**b**

| Genus | Virus name | Genome size (nt) | N* (nt) | P* (nt) | M* (nt) | F* (nt) | RBP* (nt) | L* (nt) |
|---|---|---|---|---|---|---|---|---|
| *Morbillivirus* | MBaMV | 15,804 | 1,572 | 1,524 | 1,011 | 1,995 | 1,824 | 6,552 |
| | | | (523aa) | (507aa) | (336aa) | (664aa) | (607aa) | (2,183aa) |
| | CDV | 15,690 | 1,572 | 1,524 | 1,008 | 1,992 | 1,824 | 6,555 |
| | MeV | 15,894 | 1,578 | 1,524 | 1,008 | 1,653 | 1,854 | 6,552 |
| | PDV | 15,696 | 1,617 | 1,524 | 1,008 | 1,896 | 1,824 | 6,555 |
| | PPRV | 15,948 | 1,578 | 1,530 | 1,008 | 1,641 | 1,830 | 6,552 |
| | RPV | 15,882 | 1,578 | 1,524 | 1,008 | 2,250 | 1,830 | 6,552 |
| | CeMV | 15,702 | 1,572 | 1,521 | 1,008 | 1,659 | 1,815 | 6,552 |
| | FeMV | 16,050 | 1,560 | 1,476 | 1,014 | 1,632 | 1,788 | 6,609 |

**c**

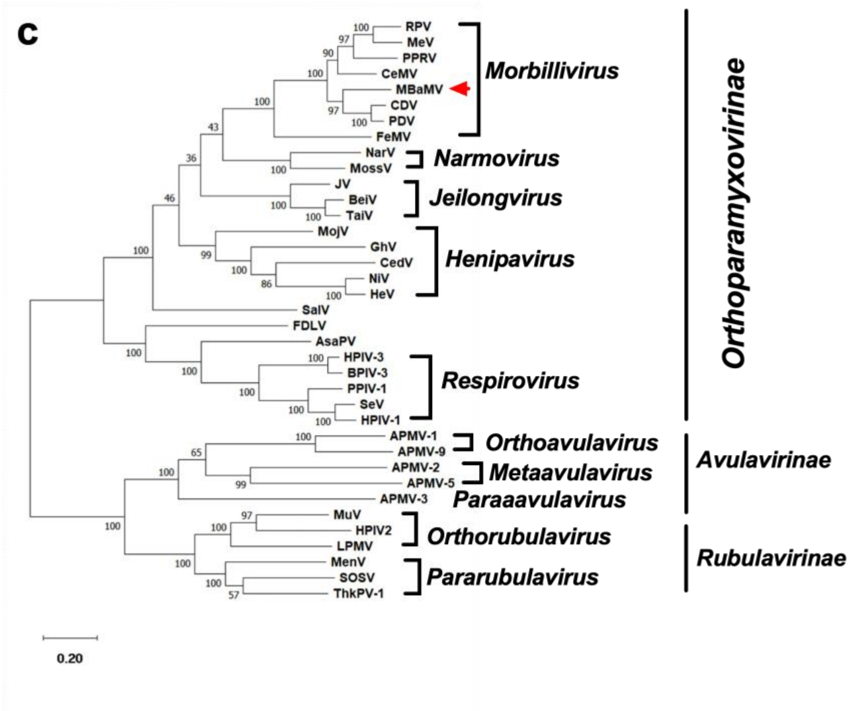


**Extended Data Figure S1. The genomic composition of myotis bat morbillivirus (MBaMV) and its phylogeny. a,** Schematic representation of the MBaMV genome and its encoded genes. The recombinant MBaMV-eGFP generated in this study is also shown. eGFP transcriptional unit is depicted as 'G' (green box). # indicates the position in L where silent mutations were introduced to prevent expression of putative ORF-X that appeared toxic in bacteria (Extended Data Figure 3). The nucleotide scale bar is provided for context. **b,** shows the genome size and the six canonical ORFs of MBaMV in comparison with other morbilliviruses. MBaMV has a total genome size of 15,804 nucleotides (nt), comprising of N (1,572 nt), P (1,524 nt), M (1,011 nt), F (1,995 nt), RBP (1,824 nt), L (6,552 nt) genes. The number of amino acids (aa) that make up each ORF are indicated in parenthesis. * nt count includes the stop codon. N; nucleoprotein, P; phosphoprotein, M; matrix, F; fusion protein, RBP; receptor binding protein (formerly referred to as H for morbilliviruses), and L; large protein which contains the RNA-dependent RNA-polymerase. **c,** The entire amino acid sequence of the L protein was used for constructing the phylogenetic tree of representative viruses from the three major subfamilies of *Paramyxoviridae*. L protein sequences were first aligned by clustalw, then the phylogenetic tree was generated by the maximum likelihood method using MEGA 10 (version 10.1.8). The numbers at each node indicate the fidelity by bootstrap test (1,000 times). MBaMV (red arrowhead) was most closely related to canine distemper virus (CDV) and phocine distemper virus (PDV) until the recent discovery of porcine morbillivirus (PoMV). The scale indicates substitutions per site. The accession numbers for the sequences used in this phylogenetic analysis are shown at Table S1.

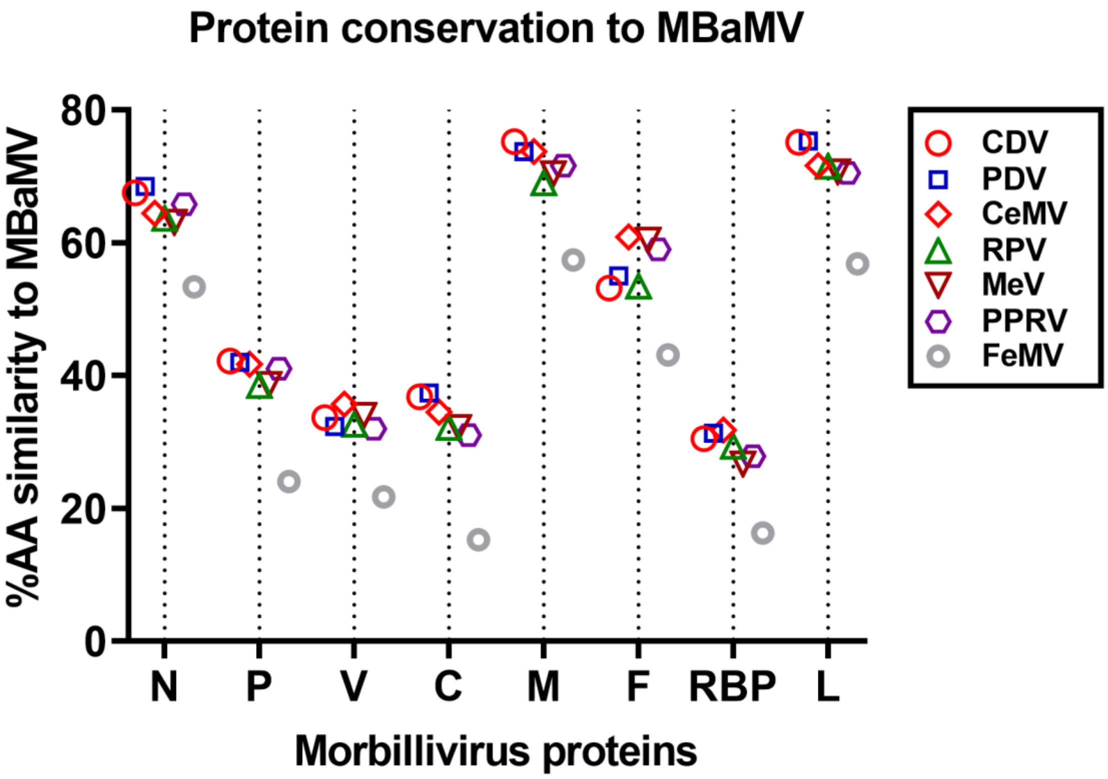


**Extended Data Figure 2. Comparison of protein amino acid sequence conservation to known morbilliviruses.** The percent amino acid similarity (% AA) of MBaMV proteins compared to the 7 established morbilliviruses are plotted to visualize the degree of conservation across the genus. N, M, F, and L showed relatively high conservation (50 – 80%) to all morbilliviruses with the exception FeMV. P, V, C and RBP showed 30-50% conservation to all morbilliviruses again with the exception of FeMV. ClustalW was used to align the cognate amino acid sequences from each morbillivirus and the % AA similarity was calculated using default settings. The accession number of the sequences used are indicated in Table S1.

a

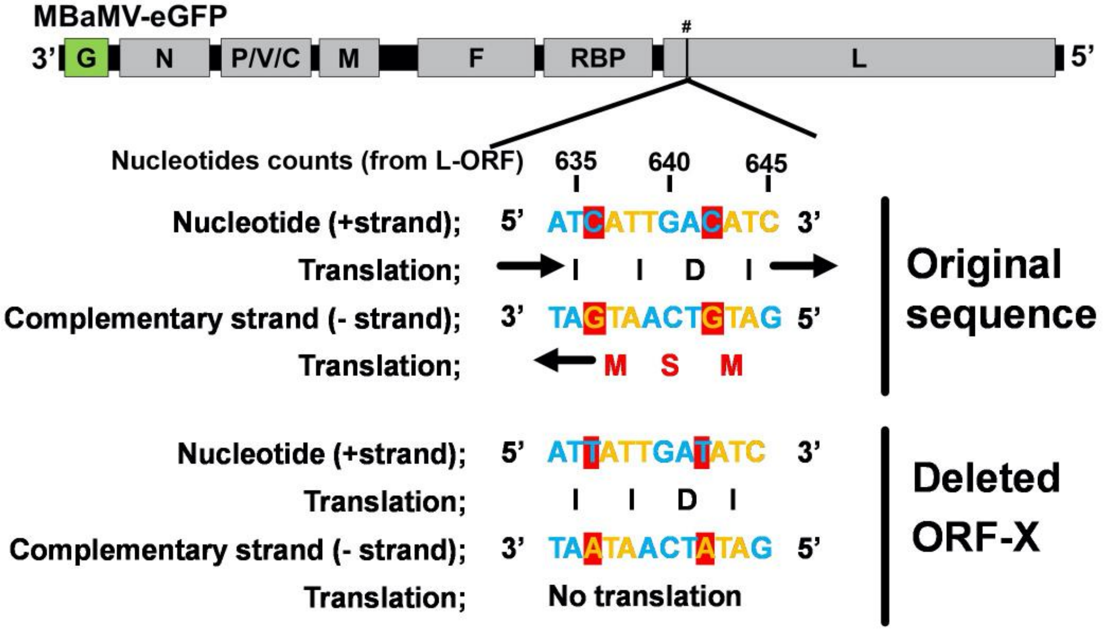


b

**Putative toxic amino acid sequence (ORF-X);**
**MSMIVTRSLVTRISYLSPLTKLGRYRRFLMLFVVVLIASLISVL**
**RVNQRRKGSNHLELHITTPKSWIIFFVVPGNPSPRPRLELT**

**Extended Data Figure 3. Deletion of putative cryptic toxic ORF in MBaMV L gene.** The first 1,500 nt of the L gene was refractory to cloning. Inspection of the L gene sequence revealed a putative ORF of 86 aa in the complementary (-) strand of the first 1,500 nt of the L gene **(a, original** sequence and **b,** ORF-X). To determine if this putative ORF might be toxic to the bacteria used for cloning the MBaMV genome, we made two mutations that were silent in the L ORF but which abolished the initiator methionine(s) in this putative ORF-X. Red shaded nucleotides correspond to third codon of methionine in ORF-X (**a,** Deleted ORF-X). This "ORF-X KO" MBaMV genomic clone was the eventual construct that was rescued. The amino acid sequence of ORF-X is shown in (**b**).

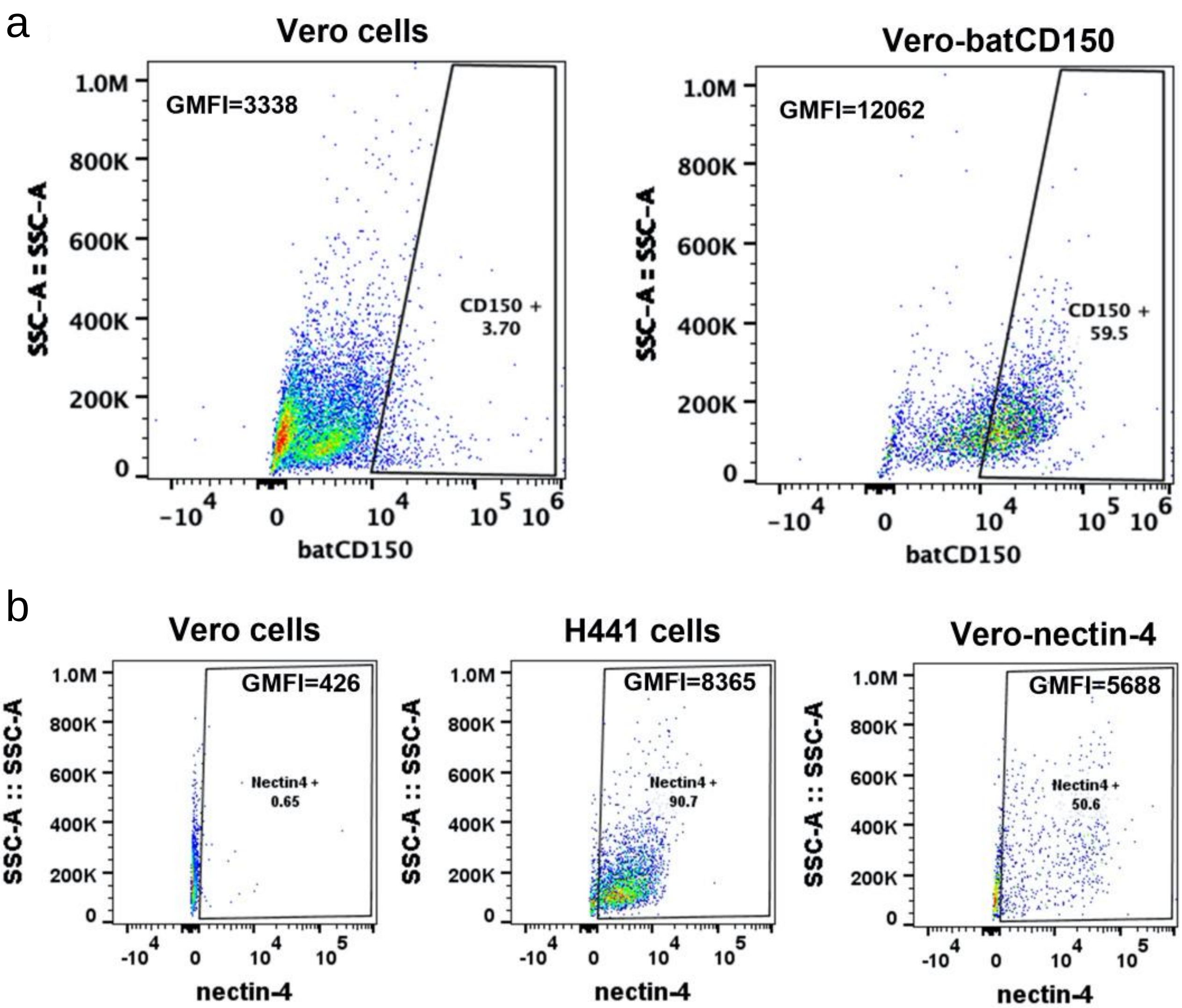


**Extended Data Figure 4. Vero cells stably expressing permissive receptors for MBaMV.** Vero cells stably expressing *Myotis brandtii* CD150 (bCD150) and human nectin-4 were made as described in methods. **a,** FACS staining for surface expression of HA-tagged bCD150 on Vero-bCD150 cells. The HA-tag was placed immediately after the signal peptide sequence. **b,** shows the human nectin-4 surface expression by FACS. Parental Vero cells and H441 (human lower airway epithelial) cells were used as negative and positive controls for human nectin-4, respectively. Greater than 50% of Vero-Nectin-4 cells express higher levels of human nectin-4 than H441 cells.

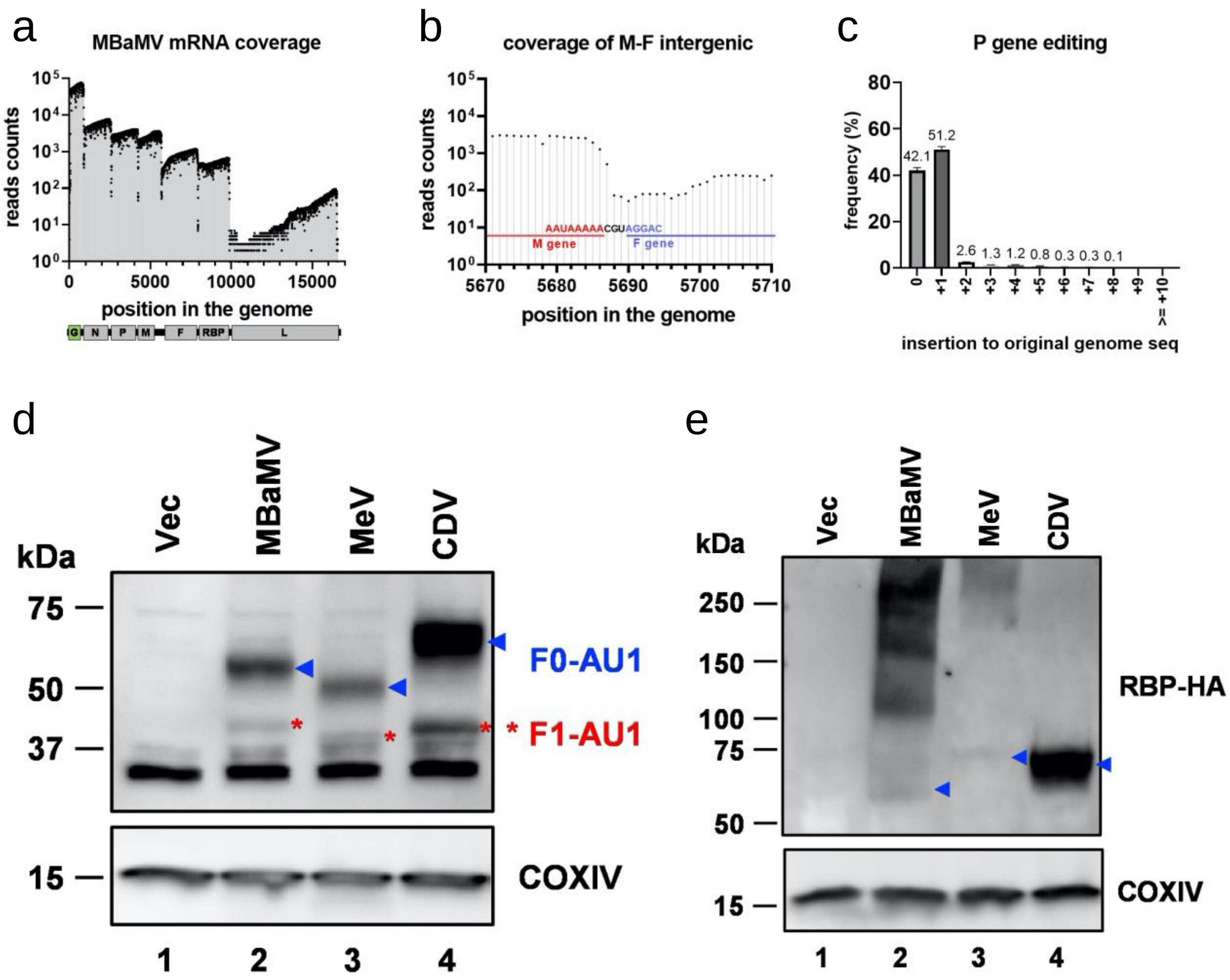


**Extended Data Figure 5. Molecular characterization of MBaMV transcripts.** Vero-bCD150 cells were infected with MBaMV (MOI=0.01) and cytosolic RNA collected at 2 dpi were subjected to direct RNA sequencing targeting mRNAs using the MinION platform (Oxford Nanopore Technology). 93,917 reads / total 797,133 reads are aligned to MBaMV. **a,** Read coverage showed that MbaMV infection and replication exhibited the 3' to 5' transcriptional gradient typical of paramyxoviruses. The boundary of each transcriptional unit can also be appreciated by the steep drop in reads that coincide with the tri-nucleotide intergenic sequence**.** Median (IQR) sequencing coverage ranges are 6509 (N, 5368-7465), 3640 (P, 3193-4047), 3018 (M, 2533-3656), 929 (F, 722-1096), 535 (RBP, 491-633) and 10 (L, 5-33). **b,** The read coverage graph around the M-F intergenic region is magnified to show the uncommon intergenic motif of 'CGU'. The intergenic motif between all other genes is 'CUU'**. c,** Amplicon sequencing around the P gene editing site shows the frequency of P gene-mRNA editing. 51.2% of P gene mRNAs acquired a single 'G' insertion at the editing site, creating V mRNA. The numbers shown are the average of three independent experiment with error bars representing S.D. (d-e) show the expression of surface glycoproteins (RBP and F) of MBaMV, MeV, and CDV by Western blot. 5 µg expression plasmids (pCAGGS, pCAGGS-RBP-HA, pCAGGS-F-AU1) were transfected into 293T in 6-well plate. Cytosolic proteins were collected and run on acrylamide gel. AU1 staining (**d**) showed full-length F protein (F0; blue arrowhead) and cleaved F1 (red asterisk) product. HA staining (**e)** showed expression of RBP monomer (blue arrowhead) around 60 – 70 kDa in addition to oligomer. COXIV was used as loading control for blot.

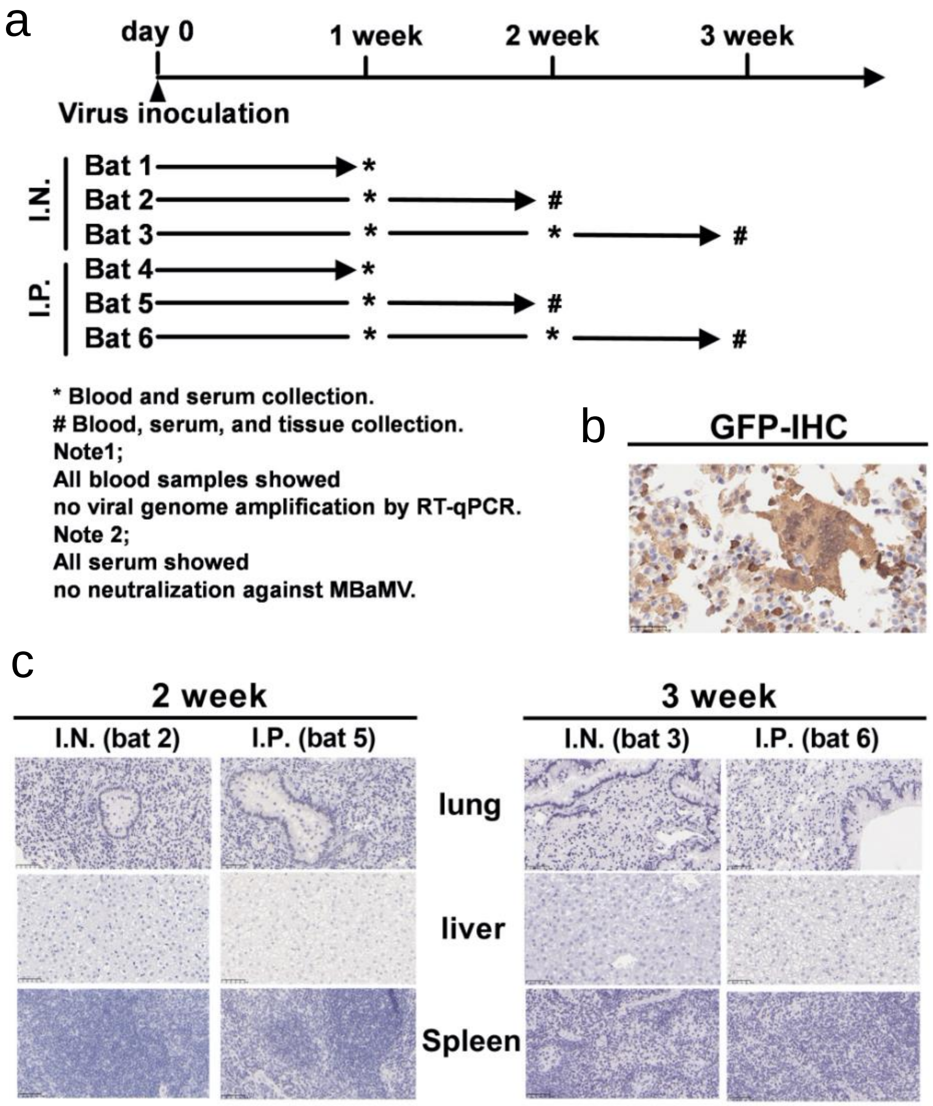


**Extended Data Figure 6. Summary of bat challenge experiment. a,** 6 Jamaican fruit bats (*Artibeus jamaicensis*) were inoculated with MBaMV (2x10$^5$ PFU/body) intranasally (I.N.) or intraperitoneally (I.P.). Blood, serum, and tissues (lung, liver, and spleen) were collected as indicated. GFP expression in the tissues were checked by GFP-**IHC. b,** MeV infected cell pellet (positive control) showed strong GFP IHC signal. **c,** bat-derived tissues showed no GFP signal.

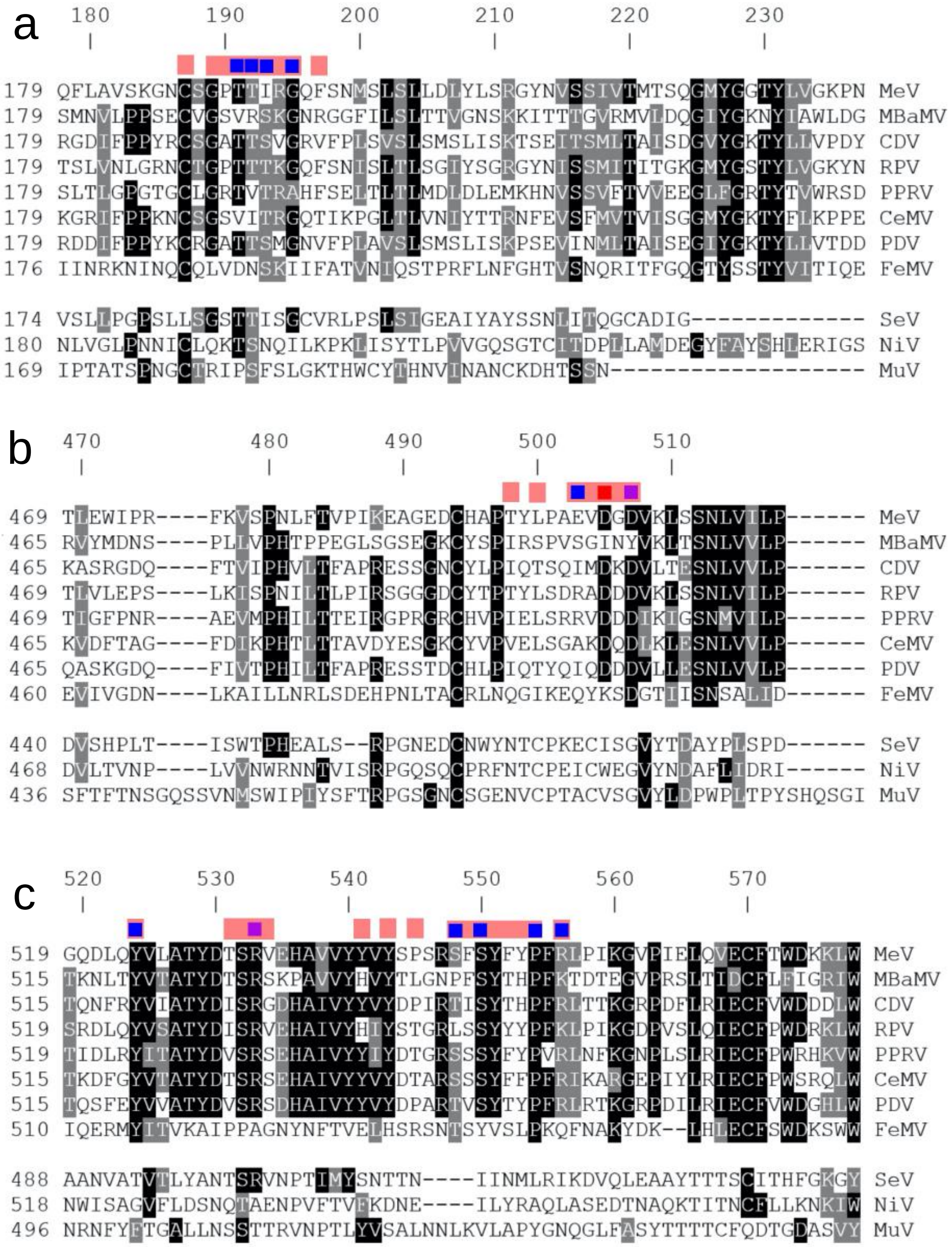

**Extended Data Figure 7. Amino acid sequence alignment of morbillivirus receptor binding proteins (RBP).** The RBP of 8 morbilliviruses and other representative paramyxoviruses (Sendai virus (SeV), Nipah virus (NiV), Mumps virus (MuV)) were aligned by clustalw. Alignments around the CD150 contact residues are shown separately in **(a)**180-200 aa, **(b)** 490-510 aa, and **(c)** 520-560 aa. Highly and moderately conserved residues in these regions are shaded in black and gray, respectively. Occluded residues at the RBP-CD150 interface from the MeV-marmoset CD150 crystal structure were identified by PDBePISA and indicated by the peach-colored shading above the alignment. Hydrogen bonds (deep blue), salt bridges (red), hydrogen bonds + salt bridge (purple) in these occluded regions are indicated.

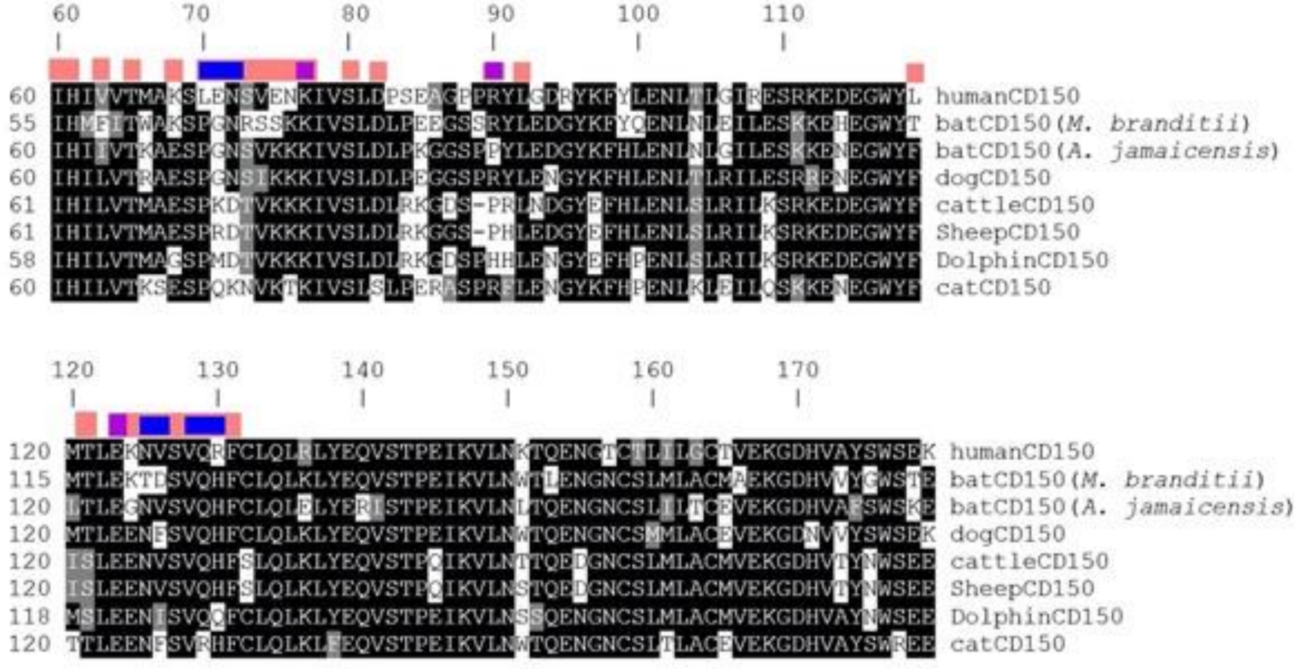

**Extended Data Figure 8. Amino acid sequence alignment of CD150 from cognate morbillivirus hosts.** CD150 from the 8 mammals that are the natural hosts of the indicated morbilliviruses are aligned by clustalw. Alignments around the RBP contact residues (aa60- 179) are shown. Highly and moderately conserved residues in these regions are shaded in black and gray, respectively. Occluded residues at the RBP-CD150 interface (PDB: 3ALZ) were identified by PDBePISA and indicated by the peach-colored shading above the alignment. Hydrogen bonds (deep blue), salt bridges (red), hydrogen bonds + salt bridge (purple) in these occluded regions are indicated. The color rendering is identical to Extended Data Figure 7.

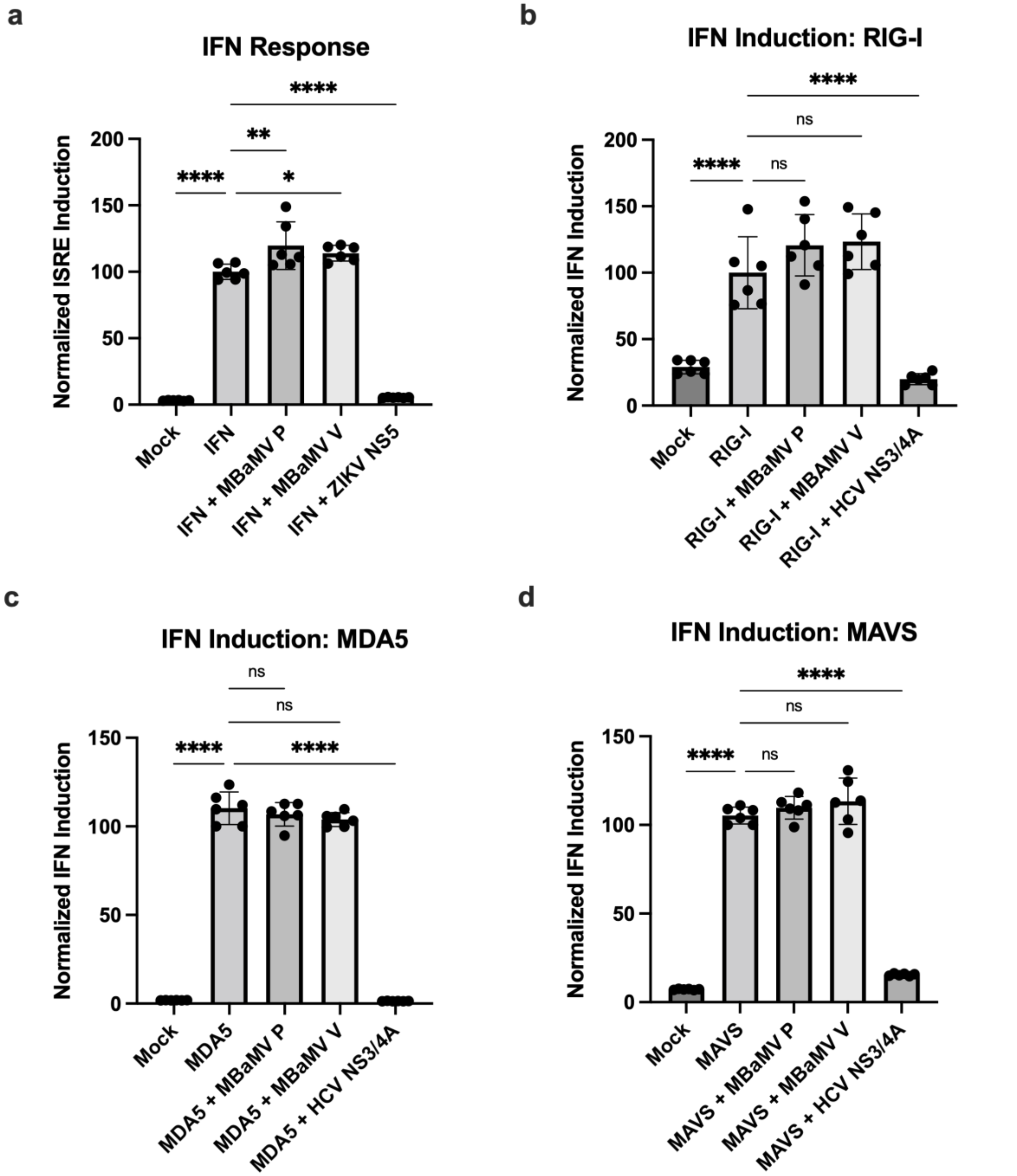


**Extended Data Figure 9. MBaMV accessory genes do not antagonize human IFN induction.** **a)** HEK 293T cells were co-transfected with an ISG54-ISRE-FLuc plasmid and either MBaMV P, MBaMV V, or ZIKV NS5. Cells were treated with human IFNb and ISRE induction was measured by luciferase assay. HEK 293T cells were co-transfected with an IFNb-FLuc plasmid, an IFN promoter stimulant **b)** RIG-I, **c)** MDA5, and **d)** MAVS, and MBaMV P, MBaMV V, or HCV NS3/4A. IFN induction was measured by luciferase assay in 2 independent experiments with 3 technical replicates.
